# Structural Basis of Substrate Recognition by the Mitochondrial ADP/ATP Transporter

**DOI:** 10.1101/2021.09.15.460093

**Authors:** Shihao Yao, Qiuzi Yi, Boyuan Ma, Xiaoting Mao, Ye Chen, Min-Xin Guan, Xiaohui Cang

## Abstract

Specific import of ADP and export of ATP by ADP/ATP carrier (AAC) across the inner mitochondrial membrane are crucial for sustainable energy supply in all eukaryotes. However, mechanism for highly specific substrate recognition in the dynamic transport process remains largely elusive. Here, unguided MD simulations of 22 microseconds in total reveal that AAC in ground c-state uses the second basic patch (K91K95R187), tyrosine ladder (Y186Y190Y194), F191 and N115 in the upper region of the cavity to specifically recognize ADP and confer selectivity for ADP over ATP. Mutations of these residues in yeast AAC2 reduce ADP transport across the *L. lactis* membrane and induce defects in OXPHOS and ATP production in yeast. Sequence analyses also suggest that AAC and other adenine nucleotide transporters use the upper region of the cavity, rather than the central binding site to discriminate their substrates. Identification of the new site unveils the unusually high substrate specificity of AAC, and together with central binding site support early biochemical findings about existence of two substrate binding sites. Our results imply that using different sites for substrate recognition and conformational transition could be a smart strategy for transporters to cope with substrate recognition problem in the highly dynamic transport process.

## Introduction

Mitochondria are the powerhouse of eukaryotic cells that synthesize ATP through oxidative phosphorylation (OXPHOS) [1,2]. The generated ATP in the mitochondrial matrix is exported to the cytosol where it is hydrolyzed to ADP to fuel cellular processes, and ADP in the cytosol is imported back to the matrix for ATP regeneration. Both ATP export and ADP import are carried out by mitochondrial ADP/ATP carrier (AAC, or ANT for adenine nucleotide translocase), the most abundant protein in inner mitochondrial membrane [3]. AAC mediates ADP/ATP exchange through alternating between cytosolic-open (c-) state and matrix-open (m-) state (Fig. 1A). The ground c-state AAC specifically recognize and import ADP, and m-state AAC specifically recognize and export ATP [4]. Matching the vital physiological significance of equal molar ADP/ATP exchange for all eukaryotic cells, AAC exhibits unusually high substrate specificity among transporters [5]. However, mechanism of the highly specific substrate recognition of AAC remains largely elusive.

**Fig. 1.**
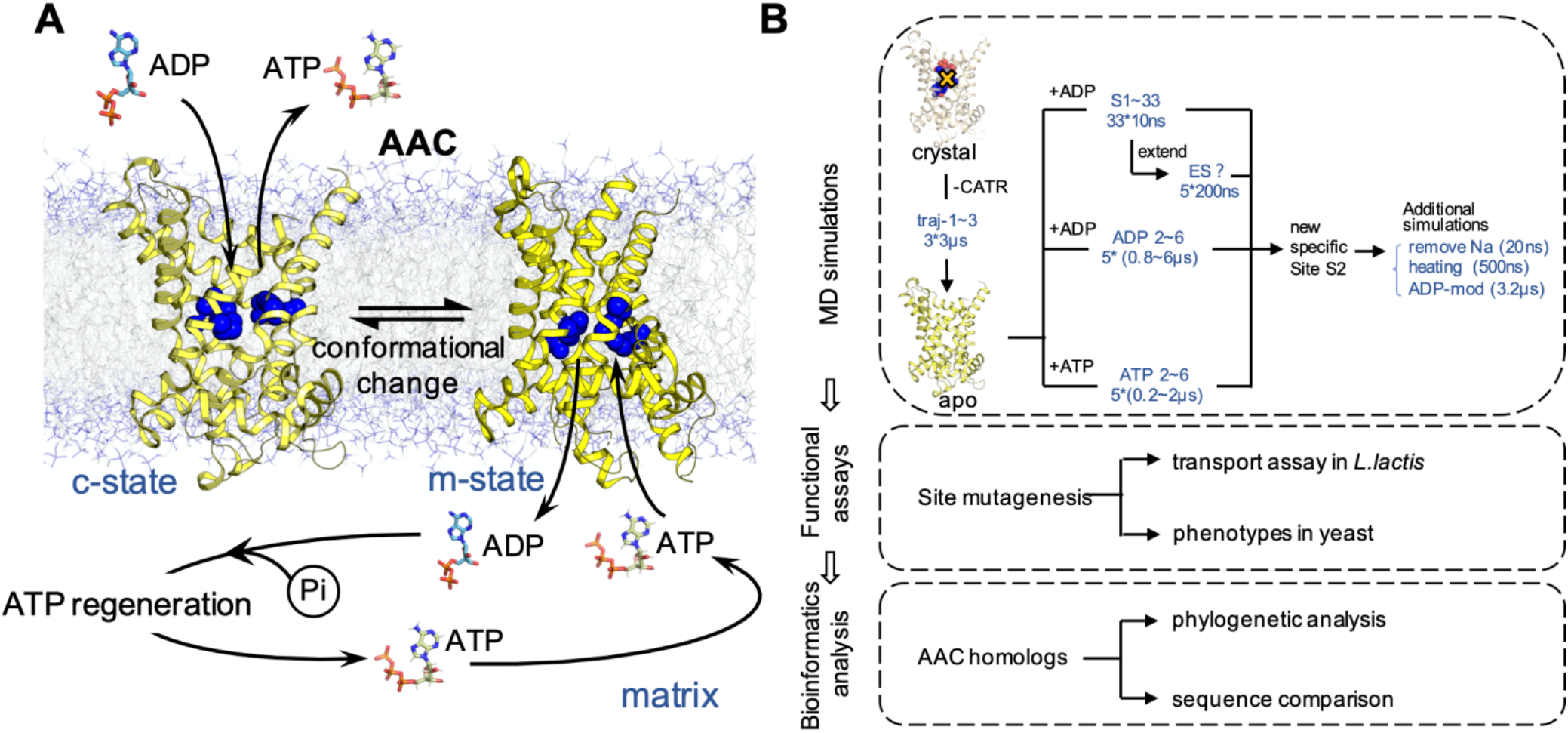
A scheme of AAC function and the workflows of this study. **(A)** A schematic diagram to show the function of AAC. The structures of AAC in c-state (PDB entry: 1OKC) and m-state (PDB entry: 6GCI) are shown. Residues of the previously proposed central binding site (K22, R79, G182, I183 and R279) are highlighted in blue spheres. **(B)** Workflow of the current work.

AAC is a paradigm of mitochondrial carrier family (MCF), the biggest solute carrier subfamily that composes of 53 members in human. MCF transporters are special for their small size and for having three homologous domains instead of two. Each domain is about 100 aa long and contains highly conserved MCF motif: Pxx[D/E]xx[K/R]xR-(20-30 residues)-[D/E]Gxxxx[Y/W/F][K/R]G [6,7], and these motif residues determine the common structural scaffold of this family. Despite similar structures of these transporters, their substrates are extremely diverse, ranging from very small proton to very large Coenzyme A (CoA) and NAD^+^. Therefore, this family are intriguing subjects to identify structural determinants for specific and selective transport which are of special significance for rational drug design.

AAC is the only MCF member whose atomic-level crystal structure has been solved up to date [8-10]. The structure of c-state AAC exhibits three-fold pseudo-symmetry, and each homologous domain consists of two TM helices connected by one short matrix helix. The six TM helices twist into a right-handed coiled coil and surround a cone-shaped transport cavity (Fig. 1A) [8]. From the crystal structure it is hard to infer the ADP binding sites because the co-crystallized inhibitor carboxyatractyloside (CATR) is much bulkier and structurally different from ADP. Analysis on the structure suggested that the featured tyrosine ladder (Y186 Y190 Y194) and four basic patches from entrance to the bottom of the cavity (K198R104, K91K95R187, K22R79R279 and K32R137R234R235 respectively) could be important in ADP binding [8,11]. Based on the crystal structure of c-state AAC, theoretical studies including molecular modeling [12], symmetry analysis [13] and molecular dynamics (MD) simulations [14,15] identified a central substrate binding site located at the bottom of the cavity. It was further proposed that this central site is the single substrate-binding site of AAC and all the other MCF members [16-18]. However, early biochemical experiments from different groups consistently demonstrated that AAC has two specific substrate binding sites: one high-affinity site and one low-affinity site [19-22]. A pair of high-affinity and low-affinity sites interacts with one CATR [23]. Moreover, the transport process was characterized as a two-step event: nucleotide binding to a specific site, followed by the vectorial process of transport [23]. Therefore, it’s apparent that more structural details, especially the structural dynamics information, are needed to explain the early biochemical findings.

Starting from the static crystal structures, all-atom MD simulations is currently the only technique to generate dynamic trajectories for proteins at atomic level. It’s especially powerful in elucidating structure-function relationship for those functionally important model molecules like AAC, for which a huge body of previous experimental results can serve as validations and provide guidance to better interpret the simulation results. Previously we used this technique to improve our understandings on the conserved MCF motif and cardiolipin binding sites of AAC [7,24]. In the current work, spontaneous ADP binding process to c-state bovine AAC1 was investigated through rigorous MD simulations of up to 22 μs in total. Improved simulation strategies enabled us to identify a new highly specific ADP binding site in the upper region of the cavity. Mutations of residues in this new ADP binding site in yeast AAC2 reduce ADP transport across the *L. lactis* membrane and induce defects in OXPHOS and ATP production in yeast. This new site was supported by sequence analysis among the five human adenine nucleotide transporters. The workflow of the current work is illustrated in Fig. 1B.

## Materials and Methods

### MD simulations

#### Large scale short MD simulations on ADP binding

Three 3-μs trajectories of *apo*-AAC were taken from our previous work [7]. Snapshots were taken from 2 μs till the end of the simulation with an interval of 100 ns. This leads to total 33 snapshots of different conformations, and for each snapshot an ADP was put near the entrance of the carrier in a random orientation and the waters or ions overlapped with the added ADP were removed. The system is composed of 70769 atoms, including one AAC molecule, 219 POPC lipids, 12226 water molecules, 23 Na^+^ and 42 Cl^-^. The GROMACS 4.5.5 package [25] is used for MD simulations, with the force field CHARMM36 [26] applied to the protein and refined CHARMM lipid parameters for the POPC molecules [27]. As each system was already fully equilibrated except the solvent around the added ADP, we ran simple equilibration steps. Two rounds of minimizations were first carried out, with and without positional constraints on the protein heavy atoms. Then each system was heated from 50 K to 310 K with positional restraints applied on Cα atoms of the protein in a time step of 2 fs, and simulation time is 50 ps. The equilibration was followed by a 10-ns production MD simulation in an NTP ensemble (310K, 1bar). The temperature of the system was maintained through the v-rescale method [28] with a coupling time of 0.1 ps, and the pressure was maintained using the Berendsen barostat [29] with τ_*p*_ of 1.0 ps and compressibility of 4.5×10^−5^ bar^-1^. The SETTLE [30] and LINCS [31] constraints were applied to the hydrogen-involved covalent bonds of water and other molecules respectively. The Particle-Mesh Ewald (PME) algorithm[32] was applied for electrostatic interactions calculation. Minimization and equilibration were carried out in local workstation carrying 32 CPUs (Intel Xeon E5-2650 2.00GHz), and production simulations were carried out on TianHe-1(A) at National Supercomputer Center in Tianjin, China. The coordinates for each system were saved every 10 ps.

#### Microsecond-timescale simulations on ADP binding and ATP binding

Five independent MD simulations were carried out on ADP binding. For comparison purpose, five simulations were also carried out on ATP binding. The ten simulations started from the same conformation of the carrier taken from the last snapshot of one trajectory in our previous work [7], but with different orientations in ADP or ATP. Other simulations protocols are the same as the short ones as described above.

#### MD simulation to promote ADP movement from the specific site S2 toward the central site S1

The simulation (ADP-mod) started from the last snapshot of ADP-3, with each atomic charge on the phosphate group of ADP reduced to 70% of normal values in the ADP topology file (the .itp file in Gromacs) and the charges of other atoms kept as the normal, and therefore the total charge of ADP was reduced to -2.3. Minimization and equilibrations were first carried out with restraints placed on both ADP and AAC, followed by a 2.1-μs production simulation. Then we continued the simulation from the check point file (.cpt file) till 3.2 μs, but used the original topology file (.tpr file) in which the charge of ADP was set to the normal values. The simulation was performed using GROMACS 4.5.5 with a time step of 2 fs. Other protocols are the same as above.

#### Trajectory analyses and free energy calculation

Most analyses were carried out with programs provided in the GROMACS package. Trajectories were viewed with VMD [33] and structural graphics were produced by the PyMOL Molecular Graphic system (version 2.0 Schrödinger, LLC). Time evolutions of the pocket volume of the c-state *apo*-AAC were calculated with POVME [34]. The line plots, edge bundling plot and free energy landscape were created in R studio with *ggplot2* and *ggraph* packages. Binding free energies were calculated with the MM/PBSA method of AMBER 14 package [35], with the GROMACS trajectories (.xtc files) converted to the AMBER format (.nc files) through the *mdconvert* command in the MDTraj toolkit and the GROMACS topology files converted to the AMBER format through ParmEd (https://github.com/ParmEd/ParmEd).

### Mutagenesis and functional assays

#### Yeast strain and cell culture media

The W301-1B (*MATα ade2-1 leu2-3,112 ura3-1 his3-22,15 trp1-1 can1-100 AAC2*) *S. cerevisiae* strain was used in this study. Media include Yeast Extract Peptone Dextrose (YPD) medium, Yeast Extract Peptone Glycerol medium and SC synthetic defined media (0.19% YNB without amino acids and NH_4_SO_4_, 2% glucose, 0.5% NH_4_SO_4_) supplemented with adenine, leucine and tryptophan (Sigma), with 1.5% agar (Sangon Biotech) added to form solid media. The media were sterilized by high pressure steam sterilization except that the glucose was treated with filter sterilization.

#### Gene knock out and mutagenesis

*AAC2* gene null strain W303-1BΔAAC2 (Δ*aac2::HIS3*) was obtained by homologous recombination[36]. The aac2 mutant alleles were obtained by overlap-extension PCR technique using appropriate primers (Table S1), with the wild type *AAC2* gene (start at -500 bp) used as template and the flag tag inserted before the end codon. The recombination fragments were digested by KpnI and BamHI (Thermo Fisher) and cloned into KpnI-BamHI-digested pFL38 plasmid by T4 ligase (NEB). The *AAC2* null strain was transformed with pFL38-AAC2-flag, pFL38-aac2-mutant-flag or the empty vector pFL38 through the Li-Ac method [37].

#### Western blot analysis

Yeast cells grew in the SC media without uracil and histidine + 2% glucose liquid medium for about 20 hours at 30°Cshaking at 220 rpm. Cells were harvested by centrifugation (4000 × g, 5 min, room temperature). Cell walls were removed by Zymolyase [38] and total cellular proteins were extracted with RIPA reagent (Invitrogen) that contains Protease Inhibitor Cocktail (Bimake). The proteins were quantitated by BCA protein determination method (Generay Biotech) before used for western blot analysis [39]. 20 mg proteins were electrophoresed through 12% bis-Tris SDS-polyacrylamide gels and then transferred to PVDF membrane. The antibodies used include anti-flag monoclonal antibody (mouse, Abcom) and anti-pgk1 Polyclonal Antibody (rabbit, Invitrogen). The secondary antibodies used include Peroxidase Affini Pure goatanti-mouse IgG and goat anti-rabbit IgG (Jackson). The ECL system (CWBIO) was used for protein signals detection.

#### Growth analysis

Monoclonal strains of yeast were picked from streak plate and amplified in the liquid screening medium as mentioned above. Cells were harvested by centrifugation, washed twice and resuspended in sterile water. The OD_600_ was measured, and the cells were diluted with sterile water to an OD_600_ of 0.5 (approximately 10000 cells/μl), based on which a ten-fold serial dilution was set up. 1 μl diluted cells were pointed on plate media of various carbon sources respectively and grown at 30°C for five days, and the growth status was recorded on daily basis [40].

#### ATP production assay

The overnight cultured yeast cells were centrifuged and resuspended in starvation medium. After 3 hours of starvation treatment, the cells were harvested and equally divided into two parts. Each part was resuspended in glucose or 2-dg/pyruvate media respectively and cultured at 30°C for 1 hour with gentle shaking. Cell walls were removed by Zymolyase before ATP assay reagent (Promega) was added. After gentle shaking for ten minutes in dark, the luminescence was measured with microplate reader Synergy™ H1(BioTek) [39].

#### Construction of Lactococcus lactis expression strains and transport Assays

Wild-type and mutant aac2 alleles were obtained by overlap extension PCR technique as mentioned before, and cloned into pNZ8048 expression vector under the control of a nisin A-induced promoter (PnisA) [41]. The plasmids carried with Chloramphenicol resistant gene were amplified in MC1061 *E*.*coli* and transformed into *L. lactis* strain NZ9800 by electroporation [42] (Lonza) which were screened on GM17 plate medium with 5 μg/ml Chloramphenicol and further determined their genotypes by sequencing.

Pre-cultures of *L. lactis* were obtained by inoculating 5 ml of GM17 medium with 5 μg/ml chloramphenicol by *L. lactis* glycerol stock from -80°C and grow at 30°C, without aeration, overnight. 50 ml of flesh GM17 medium was inoculated with pre-cultures in a dilution of 1:25 and grow at 30°C, without aeration until the OD_600_ was about 0.4. The protein was induced by addition of nisin A with a dilution of 1:3333 of spent GM17 medium from nisin A excreting *L. lactis* strain NZ9700. The cells were grown for 150 min to 180 min at 30°C, harvested by centrifugation (6000×g, 5 min, 4°C) and washed with Phosphate buffer (50mM, pH 7.0), and collected by centrifugation at 4°C [43].

Transport of [α-^32^P]-ADP (Hartmann) was initiated by the addition of 500 μL Phosphate buffer (50mM, pH 7.0) with 50 μM ADP which included 90 μCi [α-^32^P]-ADP to approximately 10 μg recombinant cells in 1.5 mL centrifuge tubes and incubated in 30°C water bath. The transport was stopped by addition of 1 mL cold Phosphate buffer at 0 min, 10 min, 15 min, 30 min, 45 min, 60 min. And the cells were rapid vacuum filtrated onto 0.45 μm filter membranes (Sangon Biotech) and washed with 5mL ice-cold Phosphate buffer for twice. The filter membranes were transfer to a scintillation vial, add 10 mL scintillation solution and measured the radioactivity retained on the membranes in liquid scintillation counter Tri-carb 4910TR (Perkin Elmer) [44].

The cell membrane proteins were extracted from the equivalent recombinant cells by ultrasonication. The cell fractions were collected by centrifugation (15000×g, 4°C) and resuspended in 80 μL PBS and employed for western blot analysis which has mentioned before. The relative quantification of AAC2 protein was carried out by comparing the gray value of the mutant bands with wild-type’s in ImageJ.

### Sequence analyses among adenine nucleotide transporters

Multiple sequence alignment was carried out on amino acid sequences of all 53 human mitochondrial carriers obtained from UniProt, and then the maximum-likelihood phylogeny tree was constructed in Mega 6, with support for the nodes calculated using Bootstrap method for 500 replications. For each of the carriers (AAC, SLC25A42, GDC, ScaMCs and SLC25A41) that fall into the same clade with AAC in phylogeny analysis, multiple sequence alignments were carried out on 41 reviewed sequences of AACs, 231 SLC25A42 sequences, 185 GDC sequences, 450 ScaMC sequences and 84 SLC25A41 sequences respectively. For the carriers excluding AACs, unreviewed sequences were also included for sequence alignment, as the number of reviewed sequences is limited. Then based on the multiple sequence alignment results, the sequence logos were created through WebLogo [38].

## Results

### Large-scale short MD simulations reveal initial ADP binding steps to c-state AAC

We first carried out large-scale short MD simulations (S1∼S33, each 10 ns long) to investigate initial ADP binding steps to the ground c-state AAC. To reduce bias, initial conformations of the carrier were picked evenly from the last 1 μs trajectories in our previous three 3-μs simulations on *apo*-AAC [7], and ADP was placed in a random orientation above the pocket entrance. The results show that in 30 out of 33 simulations, ADP was quickly (within 1 ns on average) attracted to the positive residues in the upper region of the cavity, with the highest first-contact frequencies observed on residues K205, K198, K95 and K91. Once contact, none of ADP left the carrier or moved further toward the bottom of the cavity, but the bound ADP moved dynamically among these positive residues and two major moving trends are observed: K91/K95→K91K95→K91K95R187 and K198/K205→K198K205 (Fig. 2A and Table S2). Calculations over all the 33 trajectories also show that K198K205 and the second basic patch K91K95R187 exhibit the highest ADP binding occupancies (Fig. 2B). K198R104 was reported as the first basic patch based on the crystal structure, while in our simulations R104 forms a very strong salt bridge with D195 and does not get involved in ADP binding (Fig. 2B). Therefore, our results suggest the first basic patch is composed of K198K205 instead of K198R104. Based on the simulations, ADP bound to the first basic patch is more dynamic than ADP bound to the second basic patch, and they exhibit a trend of moving from the first patch toward the second patch (Fig. 2A). To confirm this, we picked five simulations in which the ADP phosphate moiety binds to the first basic patch at the end of the 10-ns simulations (S1, S15, S24, S29, S33) and extended each of them till 200 ns. The results support our speculations: ADP in all the five extended simulations left the first basic patch (at 10 ns, 105 ns, 76 ns, 176 ns and 15 ns respectively) and moved on to bind with the second basic patch (Fig. S1). The above 10-ns simulations together with 200-ns extended simulations suggest initial steps of ADP binding: when approaching from intermembrane space, ADP is first attracted by the first basic patch (K198K205) and then quickly relayed to the second basic patch (K91K95R187).

**Fig. 2.**
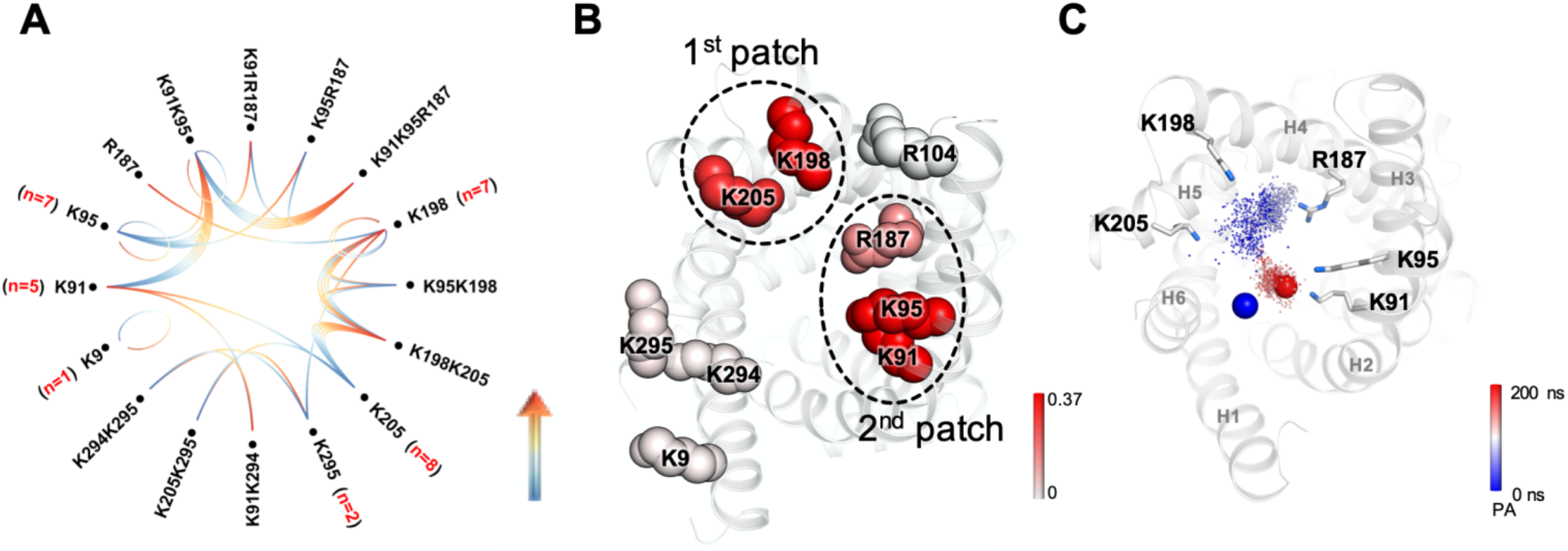
Initial ADP binding revealed by large-scale short MD simulations. **(A)** The movement of ADP phosphate moiety among the positive residues near the entrance of carrier in 31 10-ns MD simulations (in S19 and S26 ADP left the carrier in the beginning). The moving direction of each step is shown in gradient line from blue to red. The number of the simulations in which the first contact is observed is provided within the parentheses. **(B)** The ADP binding occupancies calculated on the 33 10-ns simulation trajectories are shown in colored scale on the 3D structure of c-state AAC. All positive residues are highlighted in spheres. **(C)** The trajectory of the ADP α-phosphate group in one 200-ns extended simulation (e-S33). The simulation time is shown in colored spectrum along the moving trajectory. Initial and ending positions of the ADP α-phosphate group are highlighted in blue and red spheres respectively.

### Long-time MD simulations identify a highly specific ADP binding site S2 in the upper region of the cavity

To investigate spontaneous ADP binding process on longer timescale, five long-time MD simulations (ADP-2 ∼ADP-6, ranging from 800 ns to 6 μs) were carried out, starting from the same relaxed conformation of *apo*-AAC but different orientations of ADP. Consistent with the above large-scale short MD simulations, ADP was also quickly attracted to the positive residues in the upper region of cavity and none of them left the carrier or moved further toward the bottom of the cavity. Moreover, the second basic patch also shows strong electrostatic attraction to the ADP, as judged from the long residence time (Fig. S2).

Of special significance, a highly specific ADP binding mode was consistently observed in two independent simulations (ADP-2 and ADP-3): the diphosphate moiety of ADP anchors to the second basic patch, and the adenine base forms an exquisite H-bonds network with both N115 and the backbone carbonyl oxygen of R187 (Fig. 3A). Meanwhile, the adenine base forms π-π stacking with Y190 side chain and π-cation stacking with R187 guanidinium group, leading to a sandwich stacking structure (Fig. 3A inset). In addition, 2’-OH in the ribose ring occasionally (occupancy: 9%) forms H-bond with Y190 side chain. Therefore, the five residues (K91, K95, N115, Y190 and R187) form extensive and highly specific interactions with ADP, and every part of ADP is recognized. For convenience, this newly identified specific site is denoted as S2 and the previously proposed central binding site at the bottom of the cavity is denoted as S1 hereafter in this work.

**Fig. 3.**
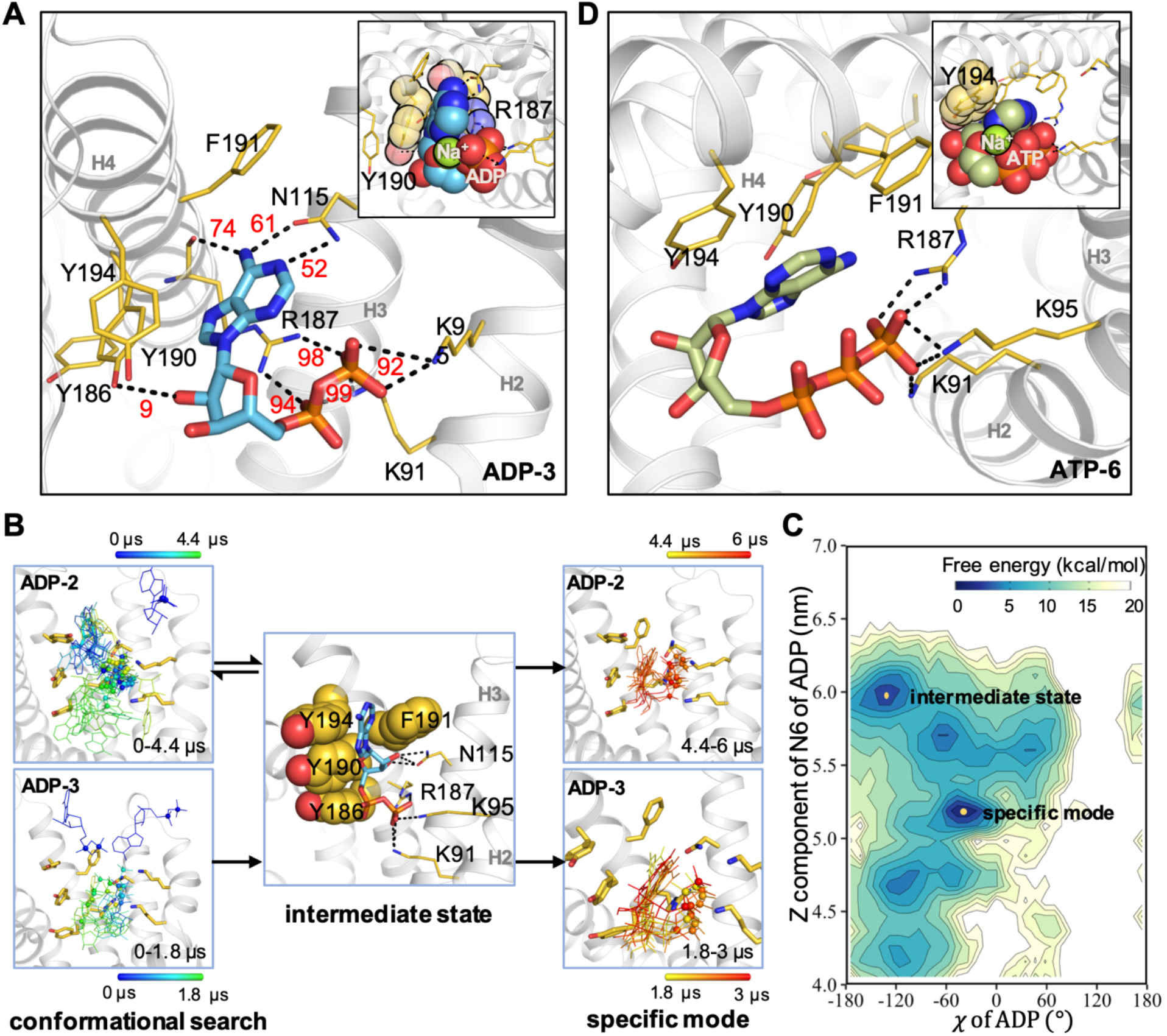
Long-time MD simulations identify a highly specific ADP binding site S2. **(A)** The highly specific ADP binding mode with site S2. Interaction occupancies are shown in red numbers. The favorable stacking structure is highlighted in the inset, with the bound sodium ion shown with a green sphere. **(B)** The consensus pathway of ADP locating to the specific binding mode at site S2 in ADP-2 and ADP-3. **(C)** Free energy landscape built from the *χ* angle and the z-component of the N6 Cartesian coordinates of ADP in ADP-2. **(D)** The non-specific ATP binding to the site S2. The favorable stacking to Y194 is highlighted in the inset, with the bound sodium ion shown in a green sphere.

Very intriguingly, ADP in ADP-2 and ADP-3 converge to the same specific binding mode with site S2 via the same pathway (Fig. 3B). The phosphate moiety of ADP was first attracted and anchored strongly to the second basic patch (Fig. S3A-C, S4A-C), while the adenine base formed dynamic stacking interactions with the tyrosine ladder (Y186 Y190 Y194), F191 and the guanidinium group of R187 (Fig. S3D, S4D). These stacking interactions help the adenine base of ADP search enormous conformations before it converges to the specific binding mode via the same intermediate conformation in the two simulations. In this intermediate conformation, ADP adopts the relaxed *anti* glycosidic conformer and the adenine base forms T-shape stacking interactions with both tyrosine ladder and F191 (Fig. 3B middle). From this intermediate conformation, ADP in the two trajectories undergoes the same changes to form the specific binding mode with site S2 (supplementary Movie S1 and Movie S2): the adenine base first experienced a rotamer change around the glycosidic bond, which leads to the unfavorable *syn* glycosidic conformation and brings the partially negative O4’ and N3 atoms to the phosphate moiety. Then a sodium ion quickly moved in to stably bind with these negative atoms. With the glycosidic conformation fixed in the *syn* conformation, ADP quickly formed the specific H-bond interactions with R187 and N115 after subtle adjustment.

In the free energy landscape built from *χ* angle and z-component of N6 coordinates of ADP, the specifically bound conformation forms the deepest basin centered at (*χ* = -47°, z(N6) = 5.13 nm) and the intermediate conformation form the second free-energy minimum centered at (*χ* = -128°, z(N6) = 6.03 nm) (Fig. 3C). A significant population of the intermediate conformation and the consistency within ADP-2 and ADP-3 suggest that the conformational changes described above might represent a predominant pathway of ADP locating to the specific mode at site S2. Although Y186, F191 and Y194 are not directly involved in specific binding with ADP (Fig. 3A), these resides are crucial in the pathway of ADP locating to the specific mode and therefore are also important part of site S2.

To demonstrate the role of the stably bound sodium ion (Fig. 3A inset), we carried out two additional simulations starting from the last snapshot of ADP-3. In one simulation we removed all sodium ions from the system, and ADP quickly (within 1 ns) lost its specifically bound conformation (data not shown). In another simulation we heated the system to 368 K, and the specifically bound sodium ion left ADP at 330 ns and then the adenine base quickly lost the specific binding mode (data not shown). These results highlight the importance of the stably bound sodium ion in maintaining the *syn* glycosidic conformation and the specific ADP binding mode at site S2.

ATP can also be imported by AAC, but at much lower efficiency. To understand mechanism of strong selectivity of c-state AAC for ADP over ATP, we also ran five MD simulations on spontaneous ATP binding (ATP-2∼ATP-6, lasting from 200 ns to 2 μs). The results show that ATP moved more dynamically among the positive residues near the cavity entrance than ADP, and ATP in ATP-3 even left the carrier at around 200 ns (Fig. S2). Intriguingly, ATP in ATP-6 binds to the site S2 in a conformation quite similar to the specifically bound ADP (Fig. 3D). The ATP also adopts the same non-canonical *syn* conformation which is stabilized by a sodium ion (Fig. 3D inset). However, because of the longer triphosphate tail, when the adenine base of ATP folds back it cannot form specific H-bonds with R187 or N115. Therefore, our simulations demonstrate sensitivity of site S2 to the length of phosphate tail of the adenine nucleotides, and ATP can only bind with site S2 in a non-specific way. MM-PBSA free energy calculations show that the ATP binding at site S2 is less favorable than ADP binding by a difference of 6 kcal/mol (Table 1), which is consistent with previous experiments showing lower affinity for ATP [39,40]. These results imply that only ADP could bind with site S2 in a highly specific mode, and c-state AAC use this site to screen different nucleotides and specifically recognize its natural substrate ADP.

**Table 1.** Binding free energy (kcal/mol) of ADP and ATP with the new site estimated by the MM-PBSA method.

| | Region used for calculation | VDWAALS | EEL | EPB | ENPOLAR | $\Delta G_{\text{gas}}$ | $\Delta G_{\text{sol}}$ | $\Delta G_{\text{bind}}$ |
| --- | --- | --- | --- | --- | --- | --- | --- | --- |
| ADP-3 | 2350-3000ns | -16.9±0.2<sup>a</sup> | -1288.9±2.2 | 1244.7±2.0 | -3.9±0.0 | -1305.8±2.2 | 1240.7±2.0 | -65.1±0.3 |
| ATP-6 | 1000-2000ns | -15.3±0.2 | -1506.6±2.3 | 1467.5±2.1 | -4.3±0.0 | -1521.9±2.3 | 1463.2±2.1 | -58.7±0.3 |

<sup>a</sup> Errors represent standard errors of mean.

During ADP conformational search, the adenine base sometimes reached very deep positions in the cavity (Fig. S5), but it did not guide ADP moving forward to the bottom of the cavity. Instead, the deeply positioned adenine base always flipped up because of the strong attachment of the diphosphate moiety to the second basic patch. This suggests that ADP will inevitably form the specific binding mode before moving forward, and to leave the site S2, the phosphate moiety of ADP has to break away from the second basic patch first.

To explore potential driving force for the specifically bound ADP to move forward, we first studied thermodynamic stability of the site S2. As shown in Fig. S6A, the whole region around site S2 is highly stable and is not affected by ADP binding to it. Potential stabilizing factors may include the short C1 loop, the strong D195:R104 salt bridge that attaches C1 loop to H4 (Fig. S6B-C) and the aromatic ladder (F90Y94F98) located at the lipid-facing interface between H2 and H3. This result suggests that ADP breaking away from the second basic patch cannot be induced by the protein local conformational changes around this site. In fact, stability of this region could be very crucial for substrate screening and specific recognition without causing obvious disturbance to the structure of the carrier. Then we did free energy calculations through MM-PBSA method. The results show that when ADP binds specifically with site S2, the binding free energy is lowered (binding affinity is increased) (Fig. S3K, S4K), while the internal energy of ADP is increased and keeps increasing at the end of the simulations (Fig. S3L, S4L). The increased internal energy is mainly caused by distorted non-canonical *syn* conformation (Fig. S3F, S4F) in which the adenine ring is almost at the same plane of the ribose ring (Fig. 3A). The above results indicate that the increased internal energy of the specifically bound ADP could be the major driving force for the ADP diphosphate moiety to break away from the second basic patch.

### An intermediate ADP conformation bound with both site S2 and the central binding site S1

The specifically bound ADP is stably maintained in our simulations although the internal energy of ADP keeps increasing before the end the simulations. Provided that one ADP/ATP exchange cycle was estimated to take 12-15 milliseconds (turnover number around 2000-2500/min at 37 °C) [41], we expect this specific ADP binding mode would last much longer than current MD simulation time scale. To speed up ADP movement in the limited simulation time, we ran a 3.2-μs simulation (ADP-mod) starting from the last snapshot of ADP-3, with partial charges of the diphosphate moiety of ADP reduced to 70% of normal values [42] before 2.1 μs and turned back to normal values after that. The simulation revealed a stable ADP conformation that binds with both site S2 and site S1 (Fig. 4A): the adenine base of ADP stacks with R187 of site S2, while the diphosphate moiety forms extensive salt bridge network with the six positive residues (K22R79R137R234R235R279) of site S1 (Fig. 4B). This conformation formed at 2342 ns and was stably maintained till the end of the simulation. Before forming this stable conformation, the diphosphate moiety of ADP first binds with K22K79 at 1311 ns and then with K22R79R137R279 at 1746 ns. During this process ADP adopts the relaxed *anti* glycosidic conformation, and the adenine base keeps stacking with R187. This stacking interaction pulled the side chain of R187 down to a position quite similar to that in m-state crystal structure (Fig. 4C). This implies that this ADP conformation could be an important intermediate conformation during c-state to m-state transition. MM-PBSA calculation shows that the ADP binding energy in this intermediate conformation (from 2350 ns till the end) is about -97.1±0.6 kcal/mol, much stronger than specific ADP binding mode at site S2. This is caused by more positive residues involved in binding with diphosphate moiety of ADP. Of special interest, drastic conformational change in the C1 loop near site S2 was observed (Fig. 4D), and we expect this was induced by the strong electrostatic interactions between diphosphate moiety of ADP and basic residues of site S1. This is similar to what happens in receptors: ligand binding induces drastic conformational changes at the other side of the transmembrane helices. For AAC, the whole region around site S2 is very stable and not affected by direct ADP binding, but is very sensitive to ADP binding to the distant site S1. We speculate that such thermodynamic feature of site S2 could be crucial for substrate selection and recognition in the highly dynamic transport process. The conformational change of C1 loop in state transition was also demonstrated by cysteine labeling experiment [43].

**Fig. 4.**
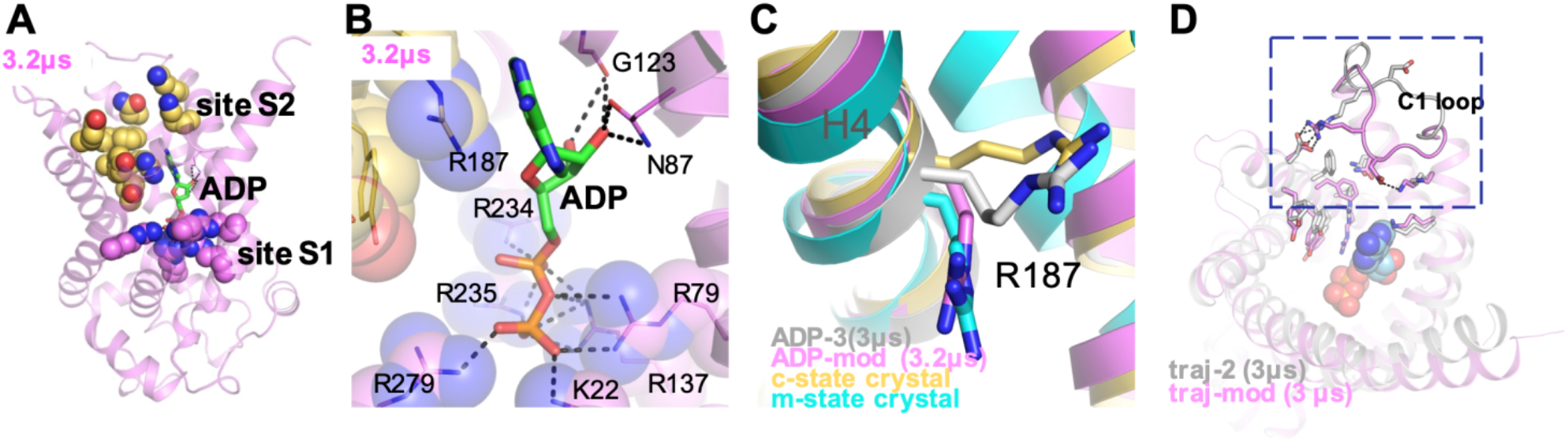
An intermediate conformation of ADP binding with the central binding site S1 and R187 of site S2. **(A)** A stable intermediate ADP conformation bridging site S2 and site S1. Residues of site S2 and site S1 are shown in yellow and violet spheres respectively, and ADP is shown in green sticks. **(B)** The salt bridge network formed between the diphosphate moiety of ADP and the positive residues of site S1. **(C)** Conformations of R187 in different structures. **(D)** Superposition of the snapshots at the end of traj-2 (silver) and ADP-mod (violet).

### Mutation of residues in site S2 of yeast AAC2 reduce ADP transport across the *L. lactis* membrane

Our MD simulations suggest that AAC uses the newly identified site S2 to specifically recognize ADP. All the eight residues of site S2 are extremely conserved among different AAC isoforms from various organisms (Fig. 5A). Moreover, these residues are highly asymmetric among three homologous domains (Fig. 5A). In fact, the second basic patch and the tyrosine ladder represent the most prominent asymmetric structural elements in the upper region of cavity in c-state AAC, matching the asymmetric structure of ADP. Yeast *AAC2* (*PET9*) is ortholog of bovine *AAC1*, and their proteins share 48% sequence identity and almost the same crystal structures (Fig. 5B). To validate functional significance of the site S2 in ADP/ATP transport, we mutated each of the eight residues to alanine in yAAC2 (Fig. 5C) and expressed the mutants on the membrane of *L. lactis* through the nisin controlled gene expression system. yAAC2 can be expressed on the membrane of *L. lactis* and can transport ADP into the cell from outside (Fig. 5D). This method was first applied to assess ADP transport activity of mutants by Kunji et al. [44]. Protein contents of AAC2 with or without mutation were first measured to verify the expression in *L. lactis* cells, and then [α-^32^P]ADP was used to measure the amount of ADP transported into the *L. lactis* cells in unit time, with the transport data normalized with the expression level of AAC2. As shown in Fig. 5E, all mutants show significant decrease in relative transport rate, with the decrease ranging from 79% to 96%.

**Fig. 5.**
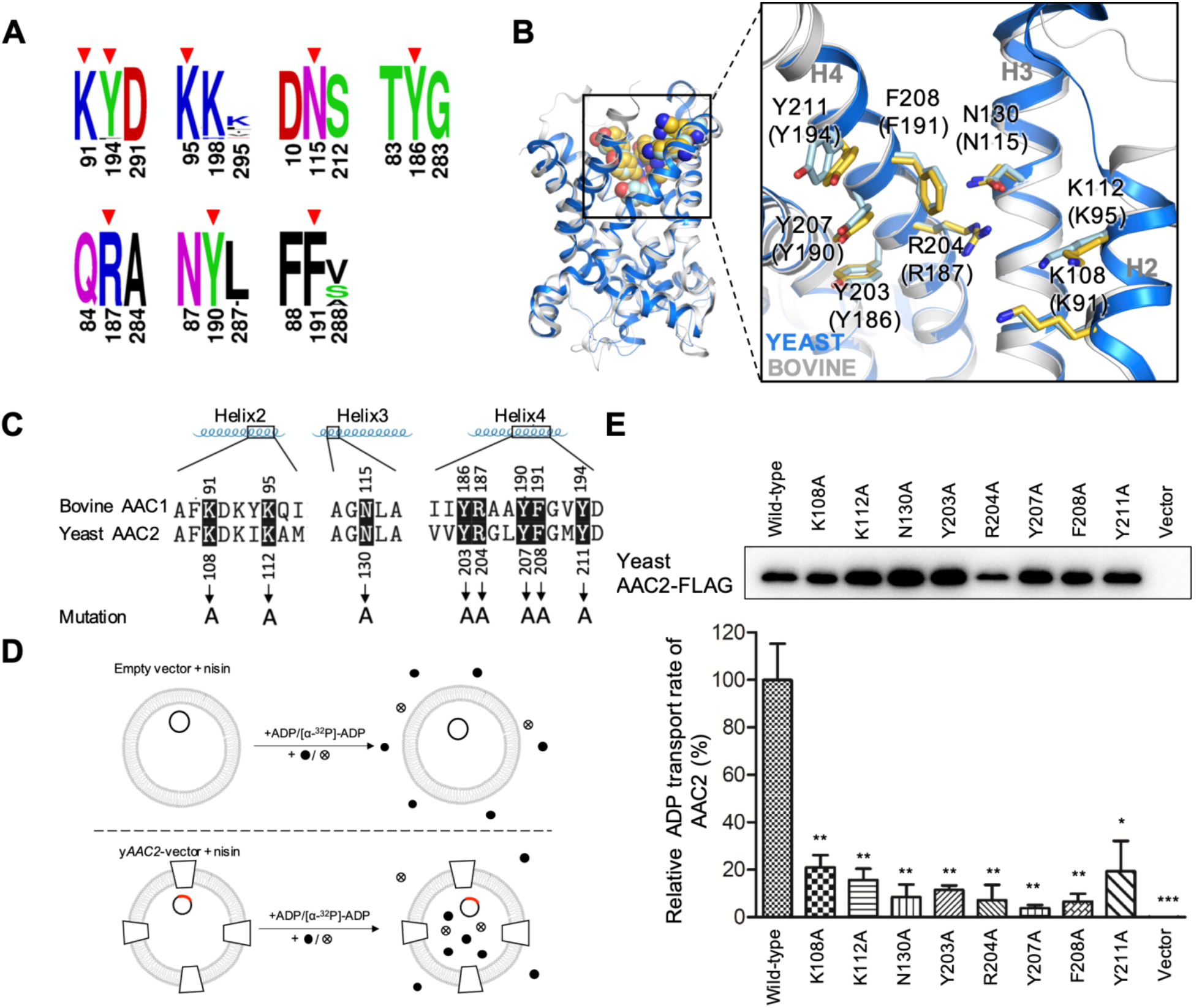
ADP transport assay across the *L. lactis* membrane. **(A)** Sequence logo presentations on the triplet to which residues of site S2 (marked with red arrows) belong. **(B)** Alignment on the structures of yeast AAC2 (PDB entry: 4C9G) and bovine AAC1 (PDB entry: 1OKC). The corresponding residues in bovine AAC1 is shown in brackets. **(C)** Sequence alignment on bovine AAC1 and yeast AAC2 and the residues chosen for alanine scanning. **(D)** The model of ADP transport assay. The black ring represents vector transformed into the *L. lactis* cells while the red part on the ring represents the *yAAC2* gene; the trapezoid represents the *yAAC2* protein on the cytomembrane; the dots in black or marked with cross represent ADP or [α-^32^P]-ADP, respectively. The lipid bilayer represents the cytomembrane of *L. lactis* cell. **(E)** The AAC2-FLAG expression levels on the membrane of *L. lactis* and the residual transport rate of the mutants relative to the wild-type measuring in whole cells of *L. lactis*. The transport rate is normalized by the expression levels of AAC2-FLAG. All Data are representative of at least three independent experiments. Data are presented as mean ± SEM, *p<0.05, **p<0.01, ***p<0.001 compared to wild-type AAC2.

### Mutation of residues in site S2 induce defects in OXPHOS and ATP production in yeast

We further did functional assays in yeast strains that express mutant yAAC2. The mutant strains were constructed by transforming *AAC2* gene with point mutation into the *AAC2*-knock out (*AAC2*-KO) yeast strain. As shown in Fig. 6A, the mutations lead to various degrees of decrease in the expression levels of mutant AAC2. The K112A and Y207A (K95 and Y190 in bAAC1) mutants showed no significant decrease (p > 0.05) in expression level compared to rescue strain, while the K108A, Y203A, R204A, F208A and Y211A (K91, N115, Y186, R187, F191 and Y194 in bAAC1) mutants showed a significantly lowered expression level.

**Fig. 6.**
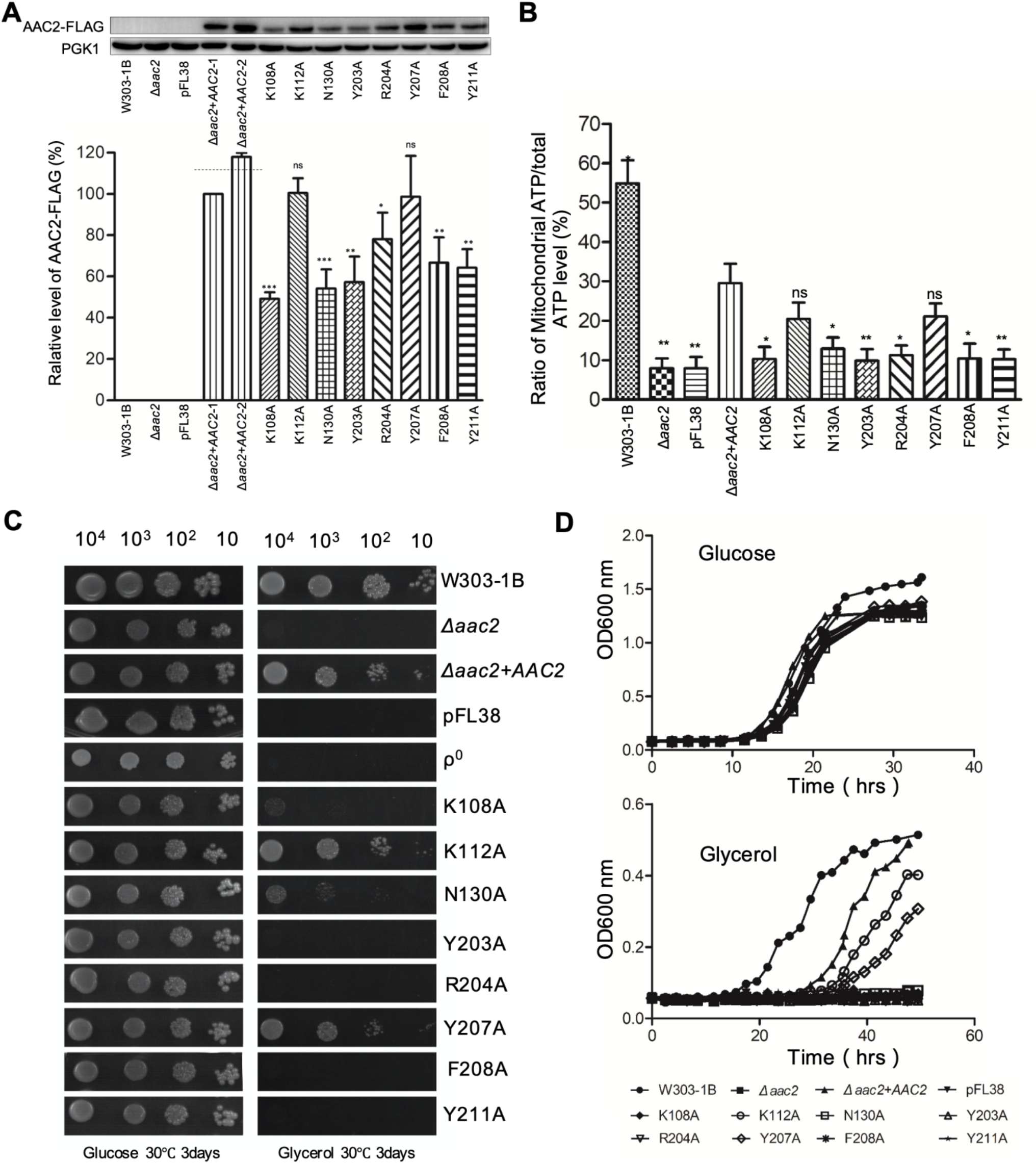
Functional assays of mutant yAAC2 in yeast. **(A)** Western blotting analysis of AAC protein expression. Twenty micrograms of total proteins from mutant and WT (W303-1B) yeast strains were electrophoresed through a denatured polyacrylamide gel, hybridized with antibodies for flag-tag as well as PGK1 as a loading control. **(B)** Ratios of mitochondrial ATP level to total ATP level in different yeast strains. **(C)** Yeast growth analysis of the wild-type and mutants on the glucose and glycerol media at 30°C. **(D)** Growth curves of yeast cells in glucose and glycerol media at 30°C. All Data are representative of at least three independent experiments. Data are presented as mean ± SEM, *p<0.05, **p<0.01, ***p<0.001 compared to rescue (Δaac2+AAC2) strain.

To assess the effect of the mutations on mitochondrial ATP production, we cultured the mutants in the glucose medium or 2-DG/Sodium Pyruvate medium and measured the ATP production respectively. As shown in Fig. 6B, the mitochondrial ATP/total ATP levels of the K112A and Y207A mutant decreased by 30%, while other mutants decreased by 56% to 67% compared to rescue strain. This data shows that the mutations lead to impaired mitochondrial ATP production.

To investigate the effect of mutations of residues in site S2 on OXPHOS, mutants were tested for growth on glucose and glycerol media. Growth of yeast using glycerol as non-fermentable carbon sources is dependent on a fully functional mitochondrion, while on glucose they can use glycolytic pathway as ATP energy source. Therefore, yeast with des-functional mitochondrion can only grow on glucose but not on glycerol. As shown in Fig. 6C, all mutants were able to grow on glucose, but showed lag in growth on glycerol. The mild lag of growth in K112A and Y207A mutants were not obvious on glycerol culture plates (Fig. 6C), but were clearly demonstrated by growth curves in glycerol medium (Fig. 6D). All the other six mutants showed severe growth defects on glycerol culture plates (Fig. 6C). These data clearly show that mutations of residues in site S2 of yAAC2 lead to impaired OXPHOS in yeast.

### Presence of specific site S2 in the upper region of the cavity is supported by sequence analysis among adenine nucleotide transporters

To validate the new specific ADP binding site S2, we compared sequences of transporters that are closely related to AAC. For those closely related transporters, residue variability in the substrate binding area confers specificity for their structurally similar yet different substrates. Therefore, specific substrate binding area could be predicted through identifying variable residues among the closely related paralogs. To this end, we first did phylogeny analysis on protein sequences of all 53 human mitochondrial carriers to identify the closely related paralogs of AAC. As shown in Fig. 7A, AACs fall into the same clade with SLC25A43, SLC25A42, GDC (Graves disease carrier protein, or SLC25A16), SCaMCs (short Ca^2+^-binding mitochondrial carrier, or Mg-ATP/Pi carrier) and SLC25A41. The substrate of SLC25A43 remains unknown yet, and this carrier is not included in analysis. Both c-state SLC25A42 and GDC specifically recognize and import coenzyme A (CoA) into the mitochondrial matrix. C-state SCaMCs and SLC24A41 specifically recognize and import Mg-ATP in exchange for intramitochondrial phosphate. ADP, CoA and Mg-ATP are all adenine nucleotides, yet quite different in size of tails and charge distributions (Fig. 7B). Therefore, their binding sites most likely show variability in the charged residues accordingly. To identify the variable charged positions in the five transporters, we first did multiple sequence alignment on the protein sequences from different organisms for each transporter (Fig. S7, S8), and then picked those variable positions that face the central cavity and have conserved charged residue in at least one of the five transporters. As shown in Fig. 7B, totally 12 variable charged positions have been identified. When mapping these variable positions on the structure of c-state AAC, ten of them are located at the upper region of the cavity, while only two (corresponding to D231 and R235 in AAC) are located at the bottom (Fig. 7C). Moreover, D231 and R235 are commonly shared by AACs, SLC25A42 and GDC (Fig. 7B), and hence they cannot be used to discriminate ADP and CoA. These sequence analyses indicate that AAC and other adenine nucleotide transporters use the upper region, rather than the previously proposed bottom region of the cavity, to discriminate and specifically recognize their substrates. This strongly supports our MD simulation results on identifying a highly specific binding site S2 in the upper region of the cavity.

**Fig. 7.**
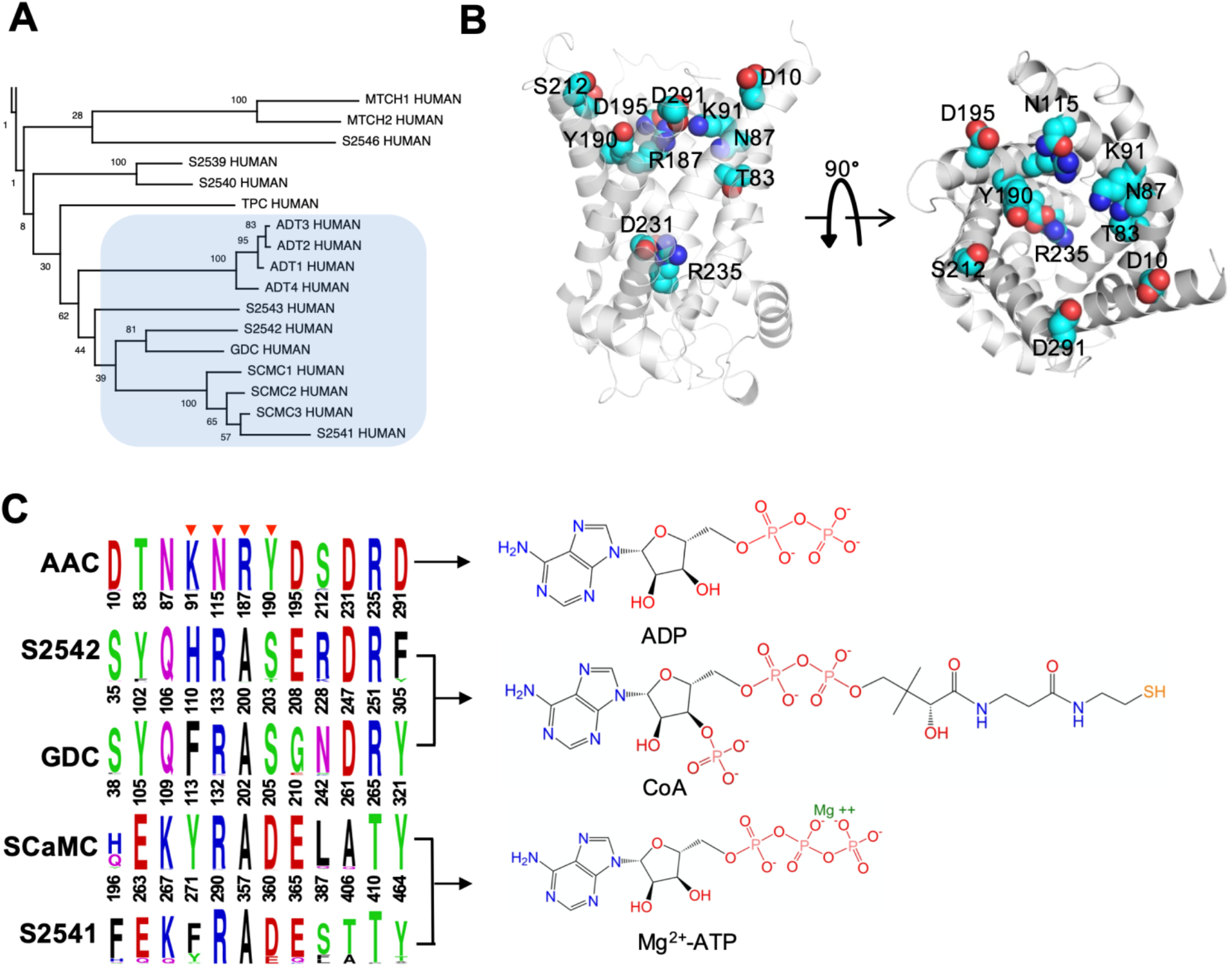
Adenine nucleotide transporters use the variable positions at the upper region of the cavity to discriminate substrates. **(A)** AACs fall into the same clade with SLC25A43, SLC25A42, GDC, SCaMCs and SLC25A41 in phylogeny analysis. **(B)** Sequence logo presentation of the variable positions among adenine nucleotide transporters and their corresponding substrates. Only variable positions that face the central cavity and involve charged residues in at least one of the five transporters are shown. Residues belongs to the specific ADP binding site S2 of AAC are highlighted with red arrows. The residues are numbered based on hAAC1, hSLC25A42, hGDC, hSCaMC1, hSLC25A41 respectively. Complete sequence logos on the odd-numbered and even-numbered helices are provided in Fig. S7, S8 respectively. **(C)** Mapping the variable positions (shown in spheres) on the c-state structure of AAC.

Among the five residues of site S2 that specifically binds with ADP, four of them (K91, N115, Y190, R187) appear at the variable positions among adenine nucleotide transporters, and these four residues are actually unique for AAC (Fig. 7B). K95 and the other three aromatic residues (Y186, F191 and Y194) of site S2 are commonly shared by adenine nucleotide transporters. Our simulations highlight the central role of R187 in specifically recognizing ADP: its side chain binds with ADP diphosphate moiety, its backbone binds with N6 atom in the adenine base of ADP, and meanwhile its side chain stacks with ADP adenine base. Actually, it forms a perfect interlocked structure with ADP, and we expect no other nucleotide could form similar structure with R187. R187 of AAC is replaced with alanine in the other four adenine nucleotide transporters (Fig. 7B). In these transporters, an arginine consistently appears at the same position equivalent to N115 of AAC, and we expect that this arginine also plays a central role in recognizing the adenine group of the substrate Mg-ATP or CoA.

## Discussion

In the current work, we’ve successfully unveiled the mechanism of unusually high substrate specificity of AAC through identifying a highly specific ADP binding site S2 that is composed of the second basic patch (K91K95R187), the tyrosine ladder (Y186Y190Y194), F191 and N115 (Fig. 3). The second basic patch and the tyrosine ladder represent the most prominent asymmetric structural elements in the upper region of the cavity in c-state AAC, matching with the highly asymmetric structure of ADP. Our MD simulations demonstrate how AAC utilizes these asymmetric structural elements to specifically recognize ADP and show selectivity over ATP. Our results are reliable in that the same highly specific ADP binding mode was consistently observed in two independent long-time MD simulations, and ADP in the two simulations converge to the specific binding mode through the same pathway and via the same highly populated intermediate ADP conformation. Mutations of residues in this new ADP binding site in yeast AAC2 reduce ADP transport across the *L. lactis* membrane (Fig. 5) and induce defects in OXPHOS and ATP production in yeast (Fig. 6). Moreover, this new site is supported by the predictions based on sequence analyses among the adenine nucleotide transporters (Fig. 7). This specific site S2 of AAC is also supported by previous NMR experiments on SCaMC, in which D360 of SCaMC, corresponding to Y190 of site S2 in AAC, was demonstrated responsible for substrate selectivity [45]. The consistency among the current work and with previous NMR experiments on SCaMC strongly suggest that the highly specific ADP binding site S2 in the upper region of the cavity in c-state AAC has been identified unambiguously.

Of special interest, five (A89D, L97P, D103G, A113P, A122D) of all the nine documented pathological mutations of AAC are located near the site S2 (Fig. S9), and previous experiments showed that these mutations lead to impaired transport of nucleotide [46]. Presence of the disease-causing mutations nearby may also serve as a justification to the newly identified binding site S2, and vice versa, identification of this new site provides potential mechanistic interpretation of these disease-causing mutations. Judging from the structure, we expect that the A122D and A89D mutations might disrupt the site S2 through forming salt bridges with R187 and K91 respectively. The L97P and A113P mutations might introduce helical backbone deformation and respectively affect K95 and N115 of site S2. The D103G mutation might destabilize the R104:D195 salt bridge that is important to stabilize the whole region around site S2.

With synthetic analog of nucleotides, previous biochemical experiments characterized the ADP transport process as a two-step event: nucleotide binding to a specific site, followed by the vectorial process of transport [23]. Moreover, conformational transition is linked to the vectorial reaction of transport, rather than substrate recognition [23]. This implies that substrate recognition and conformational transition may occur at two different sites. A bunch of early biochemical studies did show the existence of two specific ADP binding sites of different binding affinities [19-22]. These previous findings provide a framework to better interpret our simulations results, and the proposed model of ADP recognition and transport is illustrated in Fig. 8.

**Fig. 8.**
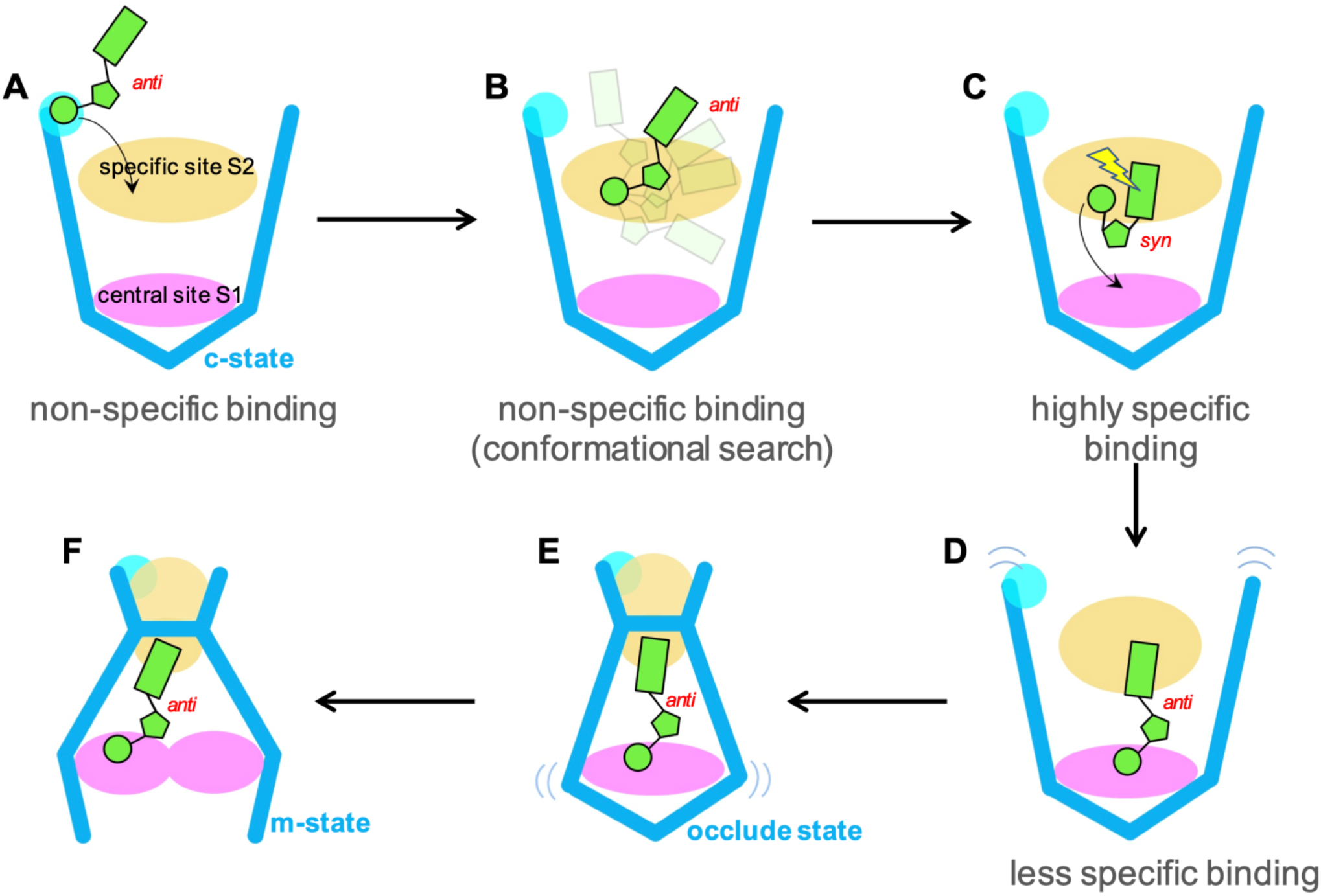
The proposed mechanism of ADP recognition and transport in AAC. ADP is shown in green, with the adenine base, ribose ring and phosphate moiety represented with rectangle, pentagon and circle respectively. The specific site S2 is shown in yellow and the central site S1 is shown in violet. The first basic patch is shown in cyan.

Our MD simulations suggest that ADP from intermembrane space are first attracted to the first basic patch (K198K205) of AAC (Fig. 8A), a conserved non-specific site that was also reported in SCaMC in previous NMR experiments [45]. Then phosphate moiety of ADP is quickly relayed to the second basic patch of the carrier, where its adenine base undergoes rigorous conformational searches favored by stacking interactions with the tyrosine ladder (Fig. 8B) before it converges to the highly specific binding mode (Fig. 8C, Fig. 3A). It’s apparent that both electrostatic interactions and stacking interactions are important in ADP recognition process. The second basic patch and the tyrosine ladder match with the negatively charged phosphate moiety and aromatic base group of nucleotides, therefore we expect that the site S2 may show non-specific attractions to a variety of nucleotides. However, only ADP could form highly specific binding mode with site S2. Significantly, the specifically bound ADP adopts the *syn* glycosidic conformation, and the increased internal energy of ADP could be the major driving force for its diphosphate moiety to break way from the second basic patch. Then the diphosphate moiety of ADP binds to the positive residues of the central binding site S1, with the adenine base still stacking with R187 of site S2. In this intermediate conformation, the strong salt bridges between the diphosphate moiety and site S1 induce drastic structural changes to the region around site S2 (Fig. 8D), which we expect will further lead to the closure of the cytoplasmic network (Fig. 8E) and then the opening of the matrix network (Fig. 8F).

In the m-state structure, F191 of site 2 together with F88 and L287 form a hydrophobic plug [17] which effectively disrupts the site S2 (Fig. S10). R187 and Y186 are located at the bottom of the cavity and are the only residues of site S2 that are accessible from the matrix side. Positive residues of site S1 are distributed in the middle region of the cavity in the m-state. Therefore, it’s possible that when AAC changes to the m-state, the adenine base of ADP may still stack with R187 and diphosphate moiety binds with part of the positive residues of site S1. That is to say, the orientation of ADP shown in Fig. 4A, B could be maintained during the c-to-m transition, with the adenine base pointing toward the intermembrane space and the phosphate moiety pointing toward the matrix. We predict that before the m-to-c transition, ATP may also adopt similar conformation: the adenine base stacks with R187, and the triphosphate moiety binds with part of the positive residues of site S1.

As mentioned above, early biochemical experiments demonstrate AAC has two specific ADP binding sites of different binding affinities [19-22]. Our calculations indicate that the newly identified site S2 corresponds to the low-affinity site, while the central binding site S1 is the high-affinity site. The current work together with previous experiments suggest that the highly specific site (S2) in the upper region of the cavity is utilized for substrate selection and recognition, while the less specific central binding site (S1) at the bottom of the cavity is mainly used for triggering conformational transition (Fig. 8). Different from receptors or enzymes, substrate binding with transporters is coupled with translocation of substrate, “movement” of binding site and more drastic conformational changes of the protein. Using the secondary binding site for substrate recognition and the central binding site only for conformational transition could be a smart strategy for transporters to cope with substrate recognition problem in the highly dynamic transport process.

## Acknowledgments

We thank Dr. Jun Ma and Dr. Ruhong Zhou for valuable advices. We thank Dr. Weifen Li (College of Animal Sciences, Zhejiang University) for Lactococcus lactis strains NZ9800/NZ9700 and pNZ8048 vector. We thank Dr. Sufen Zhang and Institute of Nuclear-agricultural science, Zhejiang University for the help on isotopic experiments. MD simulations were carried out at National Supercomputer Center in Tianjin, and the calculations were performed on TianHe-1(A).

## Author contributions

S.Y. did all the experiments, did trajectory analysis and participated in paper writing; Q.Y. carried out MD simulations, did trajectory analysis and participated in paper writing; X.M., B.M. and Y.C. did trajectory analysis and participated in paper writing; M.-X. G. supervised the project and edited the paper; X.C. conceived the project, supervised the project and wrote the paper.

