## Supplementary material for "Structural Basis of Substrate Recognition by the Mitochondrial ADP/ATP Transporter": Figures S1-10 and Tables S1-2

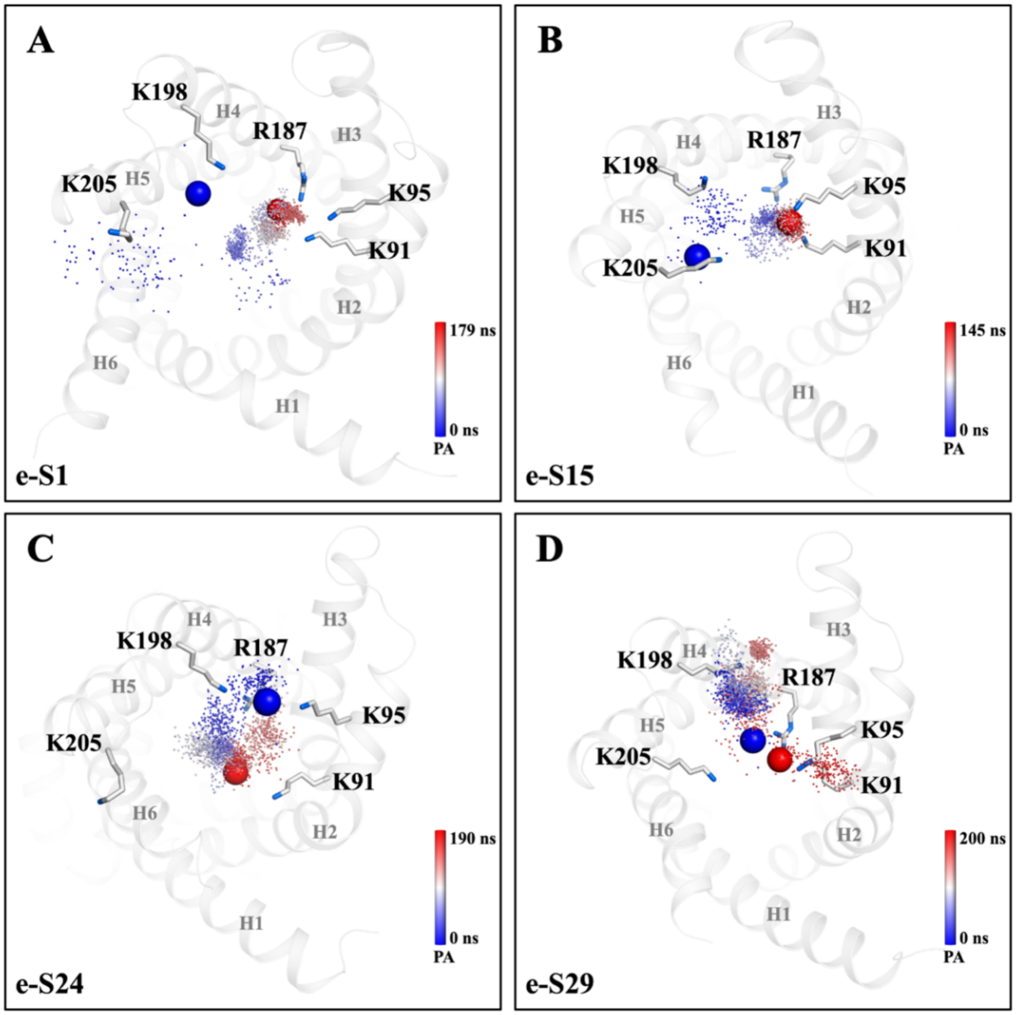


**Fig. S1 Trajectories of the α-phosphate group of ADP in the 200-ns extended simulations.** The simulation time is shown in colored spectrum, with the initial and ending positions of the ADP phosphate group highlighted with blue and red spheres respectively. The snapshots were taken every 100 ps. Residues of the first and the second basic patches are shown in sticks.

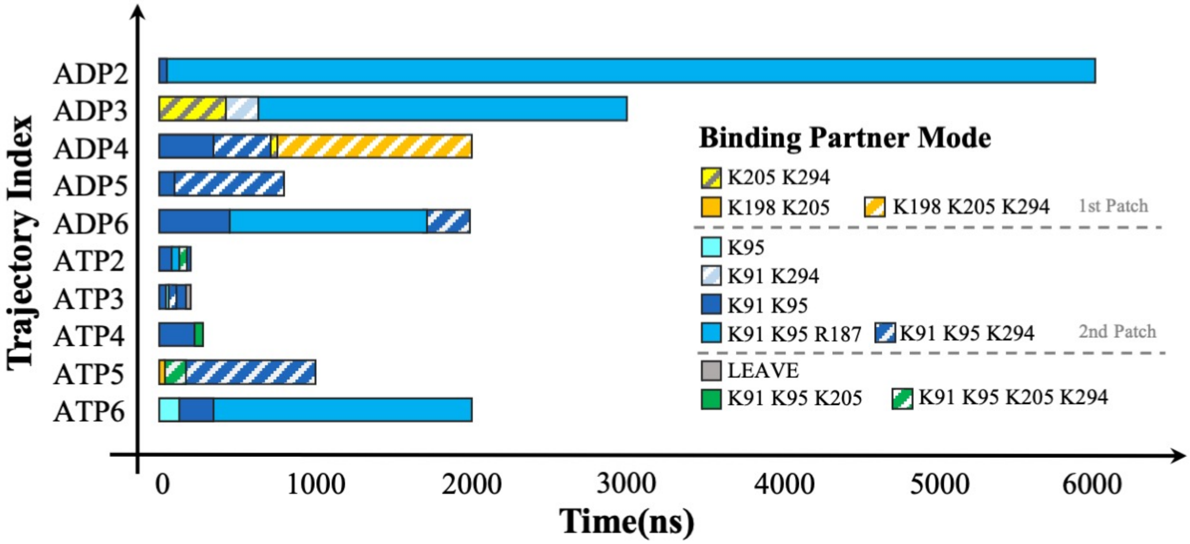


**Fig. S2 Time evolution of the bound residues with the phosphate moieties of ADP or ATP during the long time MD simulations.** More involvement of K294 in these long time simulations than in the large scale short simulations is caused by the initial conformation of AAC, in which H6 where K294 locates bends toward the center of the cavity and therefore K294 is closer to both first and second basic patches. Because this initial conformation only represents one of many possible conformations of AAC, the high occupancies of K294 in these trajectories are somewhat biased and therefore K194 is ascribed to both first and second basic patches here.

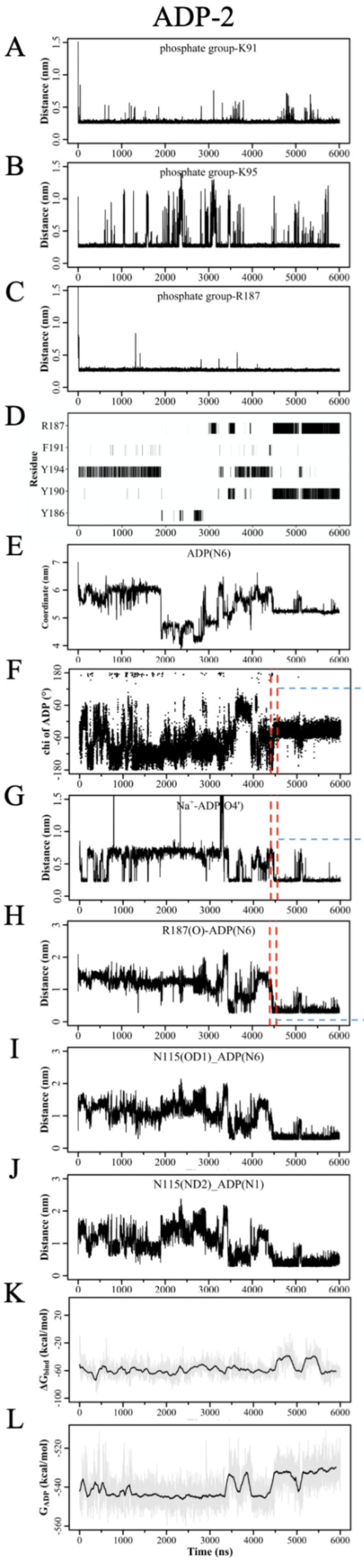


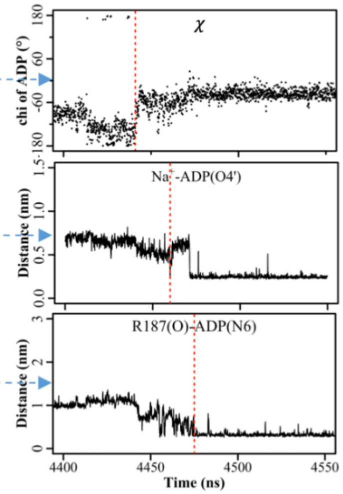


**Fig. S3 Time evolutions of the important parameters in ADP-2.** (A-C) Minimum distances between the phosphate moiety of ADP and each residue of the second basic patch. (D) π-stacking interactions with the adenine ring. (E) Z-component of the Cartesian coordinates of the N6 atom of adenine base of ADP. (F) $\chi$ angle of ADP. (G) Minimum distance between O4' atoms of ADP and sodium ions. (H) Minimum distance between N6 atom and backbone carbonyl oxygen of R187. (I, J) Minimum distances between N1 or N6 atoms of ADP and the side chain carbonyl oxygen or amine nitrogen atom of N115. (K) Binding free energy between ADP and AAC. (L) Internal energy of ADP. Some parameters around transition point are shown in detail on the right to highlight the sequence of key changes in ADP locating to the specific mode: rotamer change➔ion binding ➔H-bond formation.

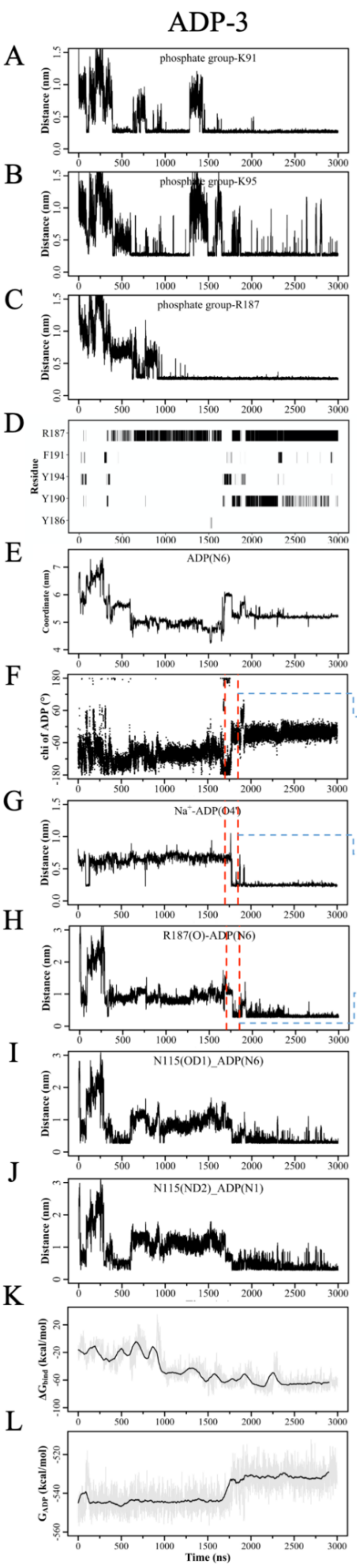


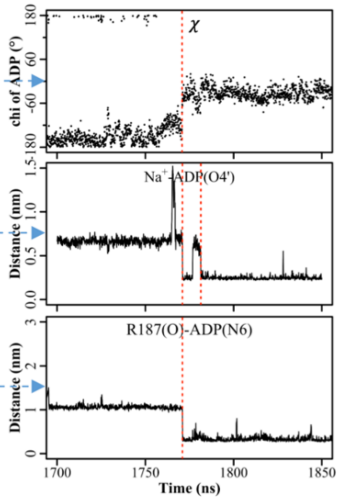


**Fig. S4 Time evolutions of the important parameters in ADP-3.** (A-C) Minimum distances between the phosphate moiety of ADP and each residue of the second basic patch. (D) $\pi$-stacking interactions with the adenine ring. (E) Z-component of the Cartesian coordinates of the N6 atom of adenine base of ADP. (F) $\chi$ angle of ADP. (G) Minimum distance between O4' atoms of ADP and sodium ions. (H) Minimum distance between N6 atom and backbone carbonyl oxygen of R187. (I, J) Minimum distances between N1 or N6 atoms of ADP and the side chain carbonyl oxygen or amine nitrogen atom of N115. (K) Binding free energy between ADP and AAC. (L) Internal energy of ADP. Some parameters around transition point are shown in detail on the right. In this trajectory, rotamer change,ion binding and specific H-bond formation occur almost at the same time.

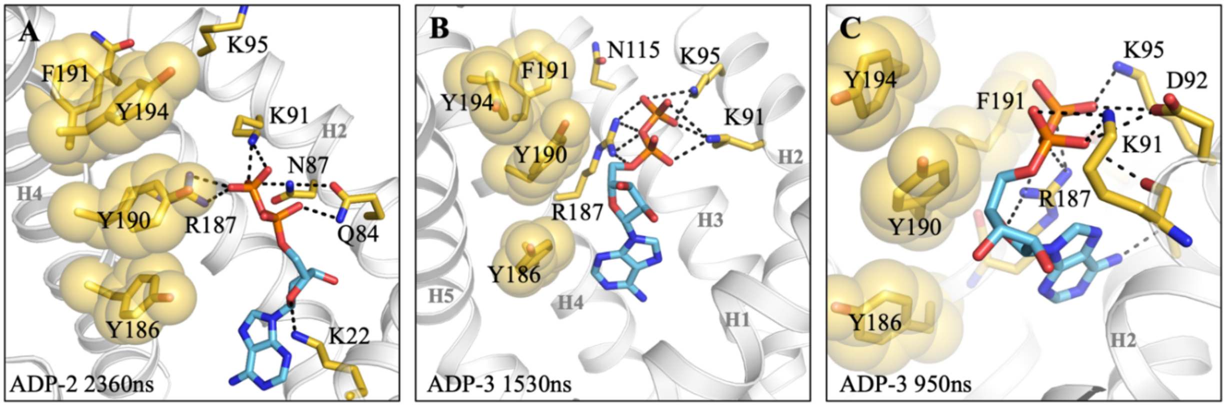


**Fig. S5 Some typical snapshots of deeply positioned adenine base of ADP at site S2.** The tyrosine ladder including F191 are shown in transparent spheres. ADP is shown in blue sticks, and the bound residues are shown in yellow sticks.

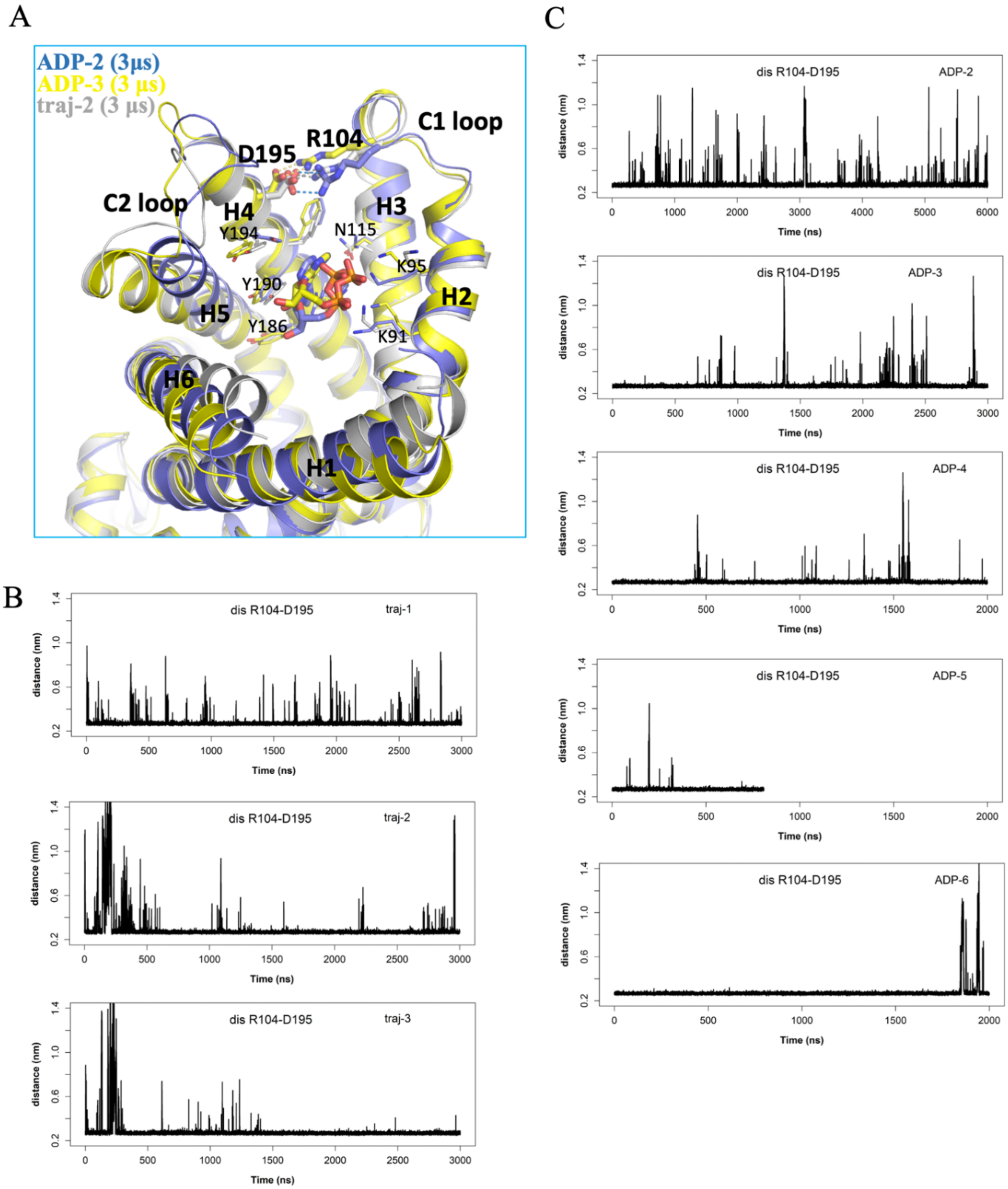


**Fig. S6 The D195:R104 salt bridge helps stabilize the region around the new specific ADP binding site.** (A) Superposition of the snapshots at the end of ADP-2 (blue), ADP-3 (yellow) and traj-2 (silver). Time evolutions of the minimum distances between the side chains of D195 and R104 in the three simulations on *apo* AAC (traj-1~3) (B) and the five long-time simulations on ADP binding (C).

A. H1

B. H3

C. H5

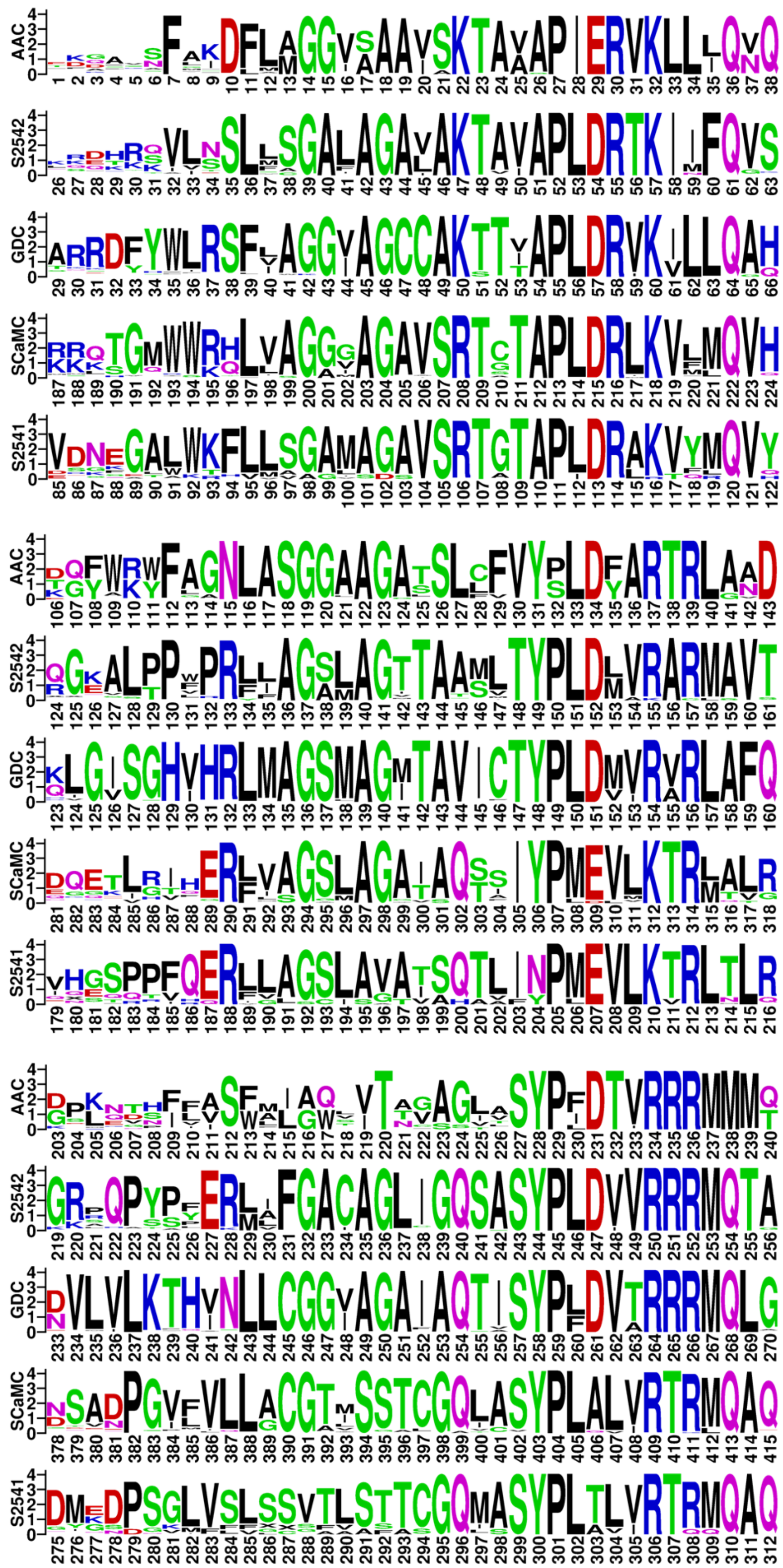


**Fig. S7 Sequence logo presentation of the odd-number helices of adenine nucleotide transporters.**

A. H2

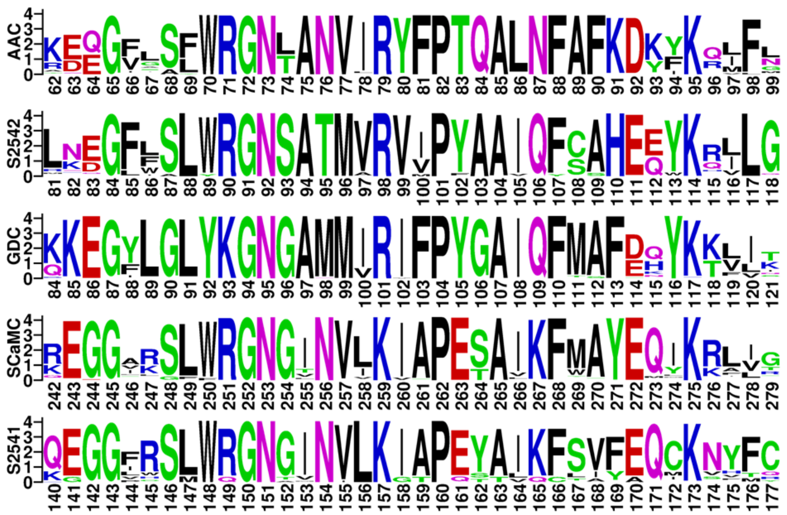

B. H4

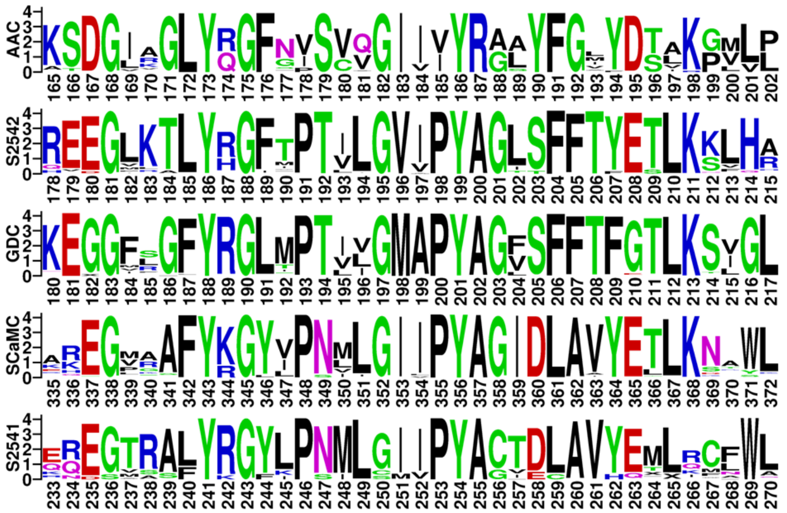

B. H6

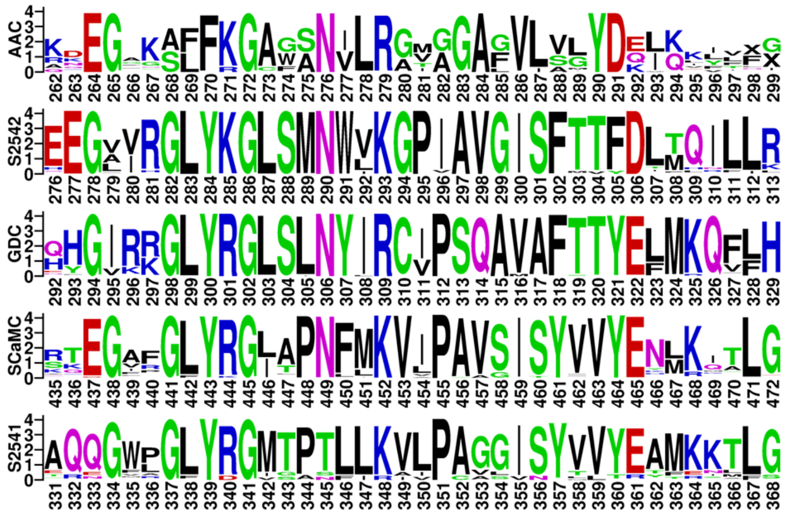

**Fig. S8 Sequence logo presentation of the even-number helices of adenine nucleotide transporters.**

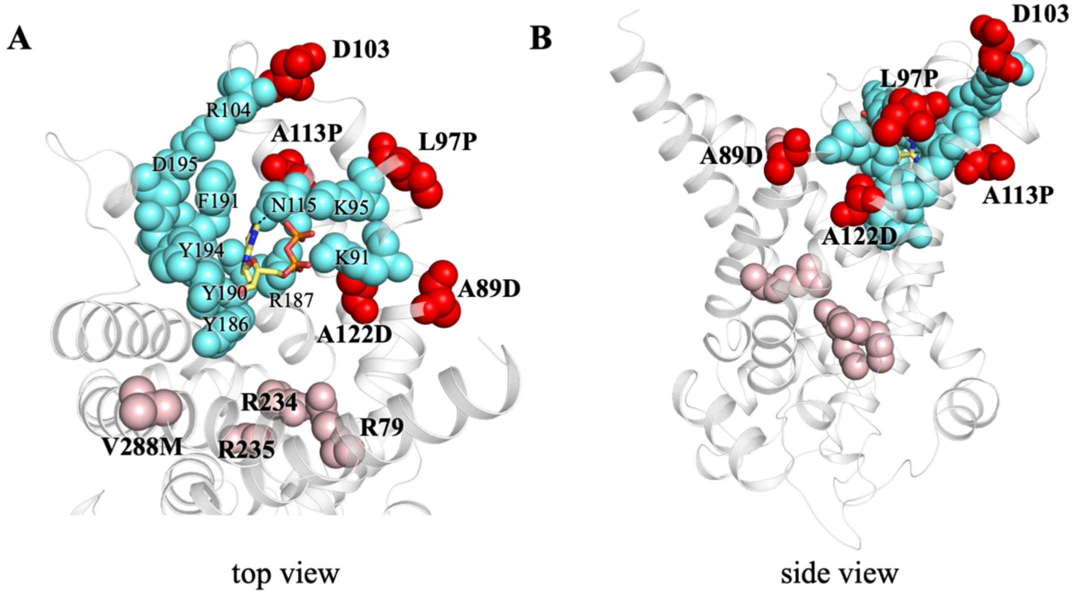


**Fig. S9 Disease-causing mutations and specific binding site S2. (A)(B)** Five documented pathological mutations (red spheres) located near the newly identified ADP specific binding site S2 (cyan spheres). Those pathological mutations not relevant to the specific site S2 are shown in pink spheres.

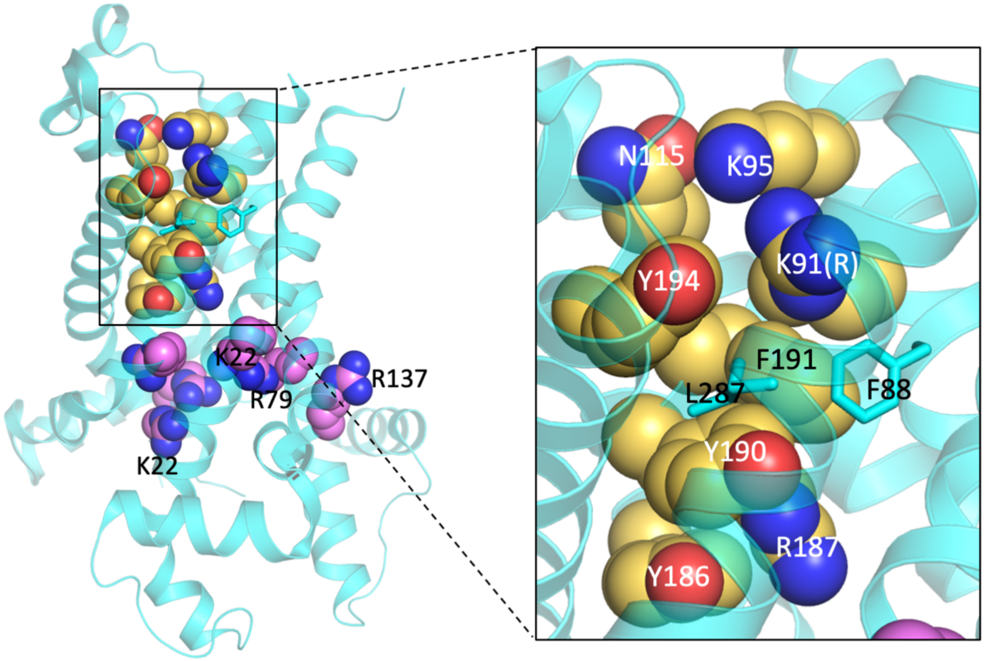


**Fig. S10 Positions of residues of site S1 and S2 in the m-state AAC.** Residues of site S1and S2 are shown in yellow and violet spheres respectively. Site S2 is disrupted by the hydrophobic plug (F88, F191 and L287) of m-state AAC and is highlighted in the inset. The structure shown here is the crystal structure of *T.thermophilus* AAC (PDB entry: 6GCI), but the residues are numbered according to bovine AAC1.

**Table S1. Oligonucleotide for AAC analysis and point mutation.**

| Primer name | Sequence |
|---|---|
| AAC2-Promoter-F-KpnI | GGGGTACCCCAGCAGCCCAACTGATAACGA |
| AAC2-FLAG-R-BamHI | CGGGATCCCGTTACTTATCGTCGTCATCCTTGTAATCTTTGAACTTC TTACCAAACA |
| Knockout-AAC2-HIS-F | ATGTCTTCTAACGCCCAAGTCAAAACCCCACTACCTCCAGCCCCA GCTCCTCTATTACTCTTGGCCTCCT |
| Knockout -AAC2-HIS-R | TTATTTGAACTTCTTACCAAACAAGATCATTTGCAGTTGGTCGTAC ATTGCCTGATGCGGTATTTTCTCC |
| MUT1-108-K-A-F | CTTTGAATTTCGCCTTCGCGGACAAGATCAAGGCC |
| MUT1-108-K-A-R | GGCCTTGATCTTGTCCGCGAAGGCGAAATTCAAAG |
| MUT2-112-K-A-F | GGACAAGATCGCGGCCATGTTTG |
| MUT2-112-K-A-R | CAAACATGGCCGCGATCTTGTCC |
| MUT3-130-N-A-F | GTTTGCCGGTGCCTTGGCATCTG |
| MUT3-130-N-A-R | CAGATGCCAAGGCACCGGCAAAC |
| MUT4-203-Y-A-F | TATTGTTGTCGCCAGAGGTCTAT |
| MUT4-203-Y-A-R | ATAGACCTCTGGCGACAACAATA |
| MUT5-204-R-A-F | TGTTGTCTACGCAGGTCTATACT |
| MUT5-204-R-A-R | AGTATAGACCTGCGTAGACAACA |
| MUT6-207-Y-A-F | CAGAGGTCTAGCCTTCGGTATGT |
| MUT6-207-Y-A-R | ACATACCGAAGGCTAGACCTCTG |
| MUT7-208-F-A-F | AGGTCTATACGCCGGTATGTACG |
| MUT7-208-F-A-R | CGTACATACCGGCGTATAGACCT |
| MUT8-211-Y-A-F | CTTCGGTATGGCCGATTCTTTGA |
| MUT8-211-Y-A-R | TCAAAGAATCGGCCATACCGAAG |

**Table S2. Movement of the diphosphate moiety of ADP among the basic residues in 33 10-ns MD simulations.**

| Traj | Movement | Traj | Movement |
|---|---|---|---|
| S1 | K205→K198K205→K198K205 | S18 | K95→K91K95→K91K95R187 |
| S2 | K198 | S19 | / |
| S3 | K205→K205K295→K295 | S20 | K295→K294K295→K91K294→K91→K91K95 |
| S4 | K91→K91K95 | S21 | K198 |
| S5 | K91R187→K91K95R187 | S22 | K95→K95R187→K91K95R187 |
| S6 | K91→K91K95 | S23 | K95→K91K95→K91K95R187 |
| S7 | K9 | S24 | K198→K198K205 |
| S8 | K95→K91K95→K91K95R187 | S25 | D103→K95 |
| S9 | K95→K91K95 | S26 | / |
| S10 | K91→K91K95 | S27 | K91K95 |
| S11 | K205→K198K205 | S28 | K198→K95K198→K198K205→K198 |
| S12 | K198→R187 | S29 | K205→K198K205 |
| S13 | K95→K91K95→K91K95R187 | S30 | K295→K198 |
| S14 | K205→K198K205→K198 | S31 | K91K95→K91R187 |
| S15 | K198K205→K95K198 | S32 | K198→K95K198→K198→K198K205 |
| S16 | K198→K198K205 | S33 | K205→K198K205 |
| S17 | K205→K91→K91K95 | | |
